# ENHANCING GENOMIC PREDICTION MODELS IN *MISCANTHUS* POPULATIONS BY INCORPORATING THE GENOTYPE-BY-ENVIRONMENT INTERACTION

**DOI:** 10.64898/2026.03.17.712492

**Authors:** Ansari Shaik, Erik Sacks, Andrew D.B. Leakey, Hua Zhao, Jens Bonderup Kjeldsen, Uffe Jørgensen, Bimal Kumar Ghimire, Alexander E. Lipka, Joyce N. Njuguna, Chang Yeon Yu, Eun Soo Seong, Ji Hye Yoo, Hironori Nagano, Kossonou G. Anzoua, Toshihiko Yamada, Pavel Chebukin, Xiaoli Jin, Lindsay V. Clark, Karen Koefoed Petersen, Junhua Peng, Andrey Sabitov, Elena Dzyubenko, Nicolay Dzyubenko, Katarzyna Glowacka, Moyses Nascimento, Ana Carolina Campana Nascimento, Maria S. Dwiyanti, Larisa Bagment, Shatabdi Proma, Julian Garcia-Abadillo, Diego Jarquin

**Affiliations:** Agronomy Department, University of Florida, Gainesville, Florida, USA; Center for Advanced Bioenergy and Bioproducts Innovation, University of Illinois at Urbana Champaign, Urbana, IL 61801, USA; Department of Crop Sciences, University of Illinois at Urbana Champaign, Urbana, IL 61801, USA; Institute for Genomic Biology, University of Illinois at Urbana Champaign, Urbana, IL 61801, USA; Department of Plant Biology, University of Illinois at Urbana Champaign, Urbana, IL 61801, USA; Center for Digital Agriculture, University of Illinois at Urbana Champaign, Urbana, IL 61801, USA; College of Plant Science and Technology, Huazhong Agricultural University, Wuhan, China; Department of Agroecology, Aarhus University, Tjele, Denmark; Bio-Herb Research Center, Kangwon National University, Chuncheon 200-701, Korea; Division of Bioresource Sciences, Kangwon National University, Chuncheon, Korea; Field Science Center for Northern Biosphere, Hokkaido University, Sapporo, Japan; FSBSI “FSC of Agricultural Biotechnology of the Far East named after A.K. Chaiki”, Ussuriisk, Russian Federation; Zhejiang Key Laboratory of Crop Germplasm Innovation and Utilization, College of Agriculture and Biotechnology, Zhejiang University, Hangzhou 310058, China; Research Scientific Computing, Seattle Children’s Research Institute, Seattle, Washington, USA; Schroll Medical ApS, Arslev, Denmark; Spring Valley Agriscience Co. Ltd., Jinan, Shandong, China; Vavilov All-Russian Institute of Plant Genetic Resources, St. Petersburg, Russian Federation; Department of Biochemistry, University of Nebraska-Lincoln, Lincoln, Nebraska; Laboratory of Intelligence Computational and Statistical Learning, Department of Statistics, Federal University of Viçosa (UFV), Viçosa, MG, Brazil; Research Faculty of Agriculture, Hokkaido University, Japan; Cand. Sci (Biology), Leading Researcher, N.I. Vavilov All-Russian Institute of Plant Genetic Resources

## Abstract

Giant *Miscanthus giganteus* (*Mxg*) is one of the most promising perennial crops to generate biomass feedstock for bioenergy and biobased products. It is derived from the natural inter-species hybridization of *Miscanthus sacchariflorus* (*Msa*) and *Miscanthus sinensis* (*Msi*) species, thus population improvement within these species is crucial. Genomic selection (GS) is an attractive option to accelerate breeding of perennial grasses, such as *Miscanthus,* which requires up to three years of evaluation to produce reliable phenotypic data. Hence, genotypes are observed in multiple years and locations causing inconsistent response patterns from one year to the next, between location, and/or location-by-year combinations. These inconsistencies are known as the genotype-by-environment interaction effect (G×E).

Although GS has been successfully implemented in multiple annual crops where straightforward cross-validation schemes exist to assess the levels of predictive ability that can be reached, for perennial crops new cross-validation schemes will help avoid data contamination. Here, we propose a series of cross-validation schemes to evaluate model performance for perennial crops. We perform a case study by analyzing one panel of each species (516 genotypes of *Msa*, 280 genotypes of *Msi*) scored for biomass yield at different locations around the world over several years. The results of the different cross-validation schemes provide insights about the usefulness of GS to accelerate the breeding process of *Miscanthus* species. In addition, leveraging the G×E effects of different types significantly increases predictive ability (up to 10% in *Msa* and 30% for *Msi*) compared to the conventional approaches based on main effects only.

## 1 INTRODUCTION

Biofuels offer a promising way to expand energy production (Saleem et al., 2022). Biomass can be used as a feedstock to make fuels such as bioethanol, biodiesel, and bio-hydrogen, as well as the energy-rich compounds needed to make many household and industrial products (Lynd & Wang, 2003; Sarkar et al., 2012). Ideal feedstock crops should produce large amounts of biomass per unit land area to maximize profits and reduce competition with other land uses. Since perennial grasses produce higher yields per unit area with fewer expensive inputs, they can be more productive and profitable as a feedstock, while requiring less land (Lewandowski et al., 2003; Heaton et al., 2008a). *Miscanthus × giganteus* is an especially promising perennial grass crop since it is a sterile, triploid hybrid (Heaton et al., 2008b; Heaton et al., 2010; Chung & Kim, 2012). The *Miscanthus* genus includes around 14-20 species closely related to sugarcane, and it stands out from other biomass crops primarily due to its large lignocellulose yields and wider adaptability (Clifton-Brown et al., 2008).

*Miscanthus* originated in tropical and subtropical regions and possesses the C4 photosynthetic pathway, which supports greater biomass production, water use efficiency and nitrogen use efficiency (Monson et al. 2025). *Miscanthus*’s wider environmental adaptability is ideal for crop establishment and distribution across various climatic conditions in Europe and North America (Numata M, 1974).

Breeding efforts are being made to develop *Miscanthus* genotypes that are highly productive as well as adapted across latitudinal and associated environmental gradients in the United States. Among all *Miscanthus* species, efforts have been focused on two species. *M. sinensis* (*Msi*), which is diploid (2n=2x=38), and *M. sacchariflorus* (*Msa*), which can be diploid or tetraploid (2n=2x=4x=76) species (Clark et al., 2014, 2015, 2019a; Clifton-Brown et al., 2008; Dwiyanti et al., 2013). These two species are of interest because they are parental lines of *M.* × *giganteus*, which is a triploid that shows wider adaptation and higher biomass production compared to the parent species. *M.* × *giganteus* for this reason, it necessitates vegetative propagation, which is costly for establishing commercial cultivation (Lewandowski et al., 2000; Chung & Kim, 2012).

*Miscanthus* is a long-lived perennial plant that requires multiple years of field testing to collect significant biomass production data (Clark et al., 2019b) to allow accurate selection. Environmental conditions influencing trait performance vary from site to site, or year to year, complicating the selection process in breeding programs (Jarquin et al., 2017). Thus, the establishment of multi-environment trials is required to ensure the stability of cultivar-specific performance, and this adds time and expense to the breeding process. Previous studies by Crossa et al. (2017) and Heslot et al. (2012) have shown that genomic selection GS enhances selection intensity by reducing the phenotyping cost of new maize breeding lines. GS employs a training population with both genotypic and phenotypic data for model calibration (Fonseca et al., 2021); then, it is used to predict the performance of candidate genotypes when only genomic information is available (Bernardo, 1994; Meuwissen et al., 2001). The main idea is to use genomic information from untested individuals (candidate population) to estimate their genomic estimated breeding values (GEBVs) and conduct the selection using these as surrogates of phenotypes. GEBVs do not explain gene functions but serve as the optimal selection criterion (Meuwissen et al., 2001; Jannink et al., 2010). GS holds the potential to increase genetic gain (van der Werf, 2013) by skipping field trials and increasing the testing capacity. Implementing GS could be helpful for selecting superior genotypes within *M. sacchariflorus* and *M. sinensis* species, enhancing their adaptability and increasing biomass production for a wider range of environmental conditions. Njuguna et al. (2023) have shown in *M. sacchariflorus* that selection methods assisted by genomic information, such as genome-wide association study (GWAS) and genomic selection (GS), can be highly successful in accelerating crop breeding. Similar conclusions were presented by Clark et al. (2019c), who focused on *M. sinensis*.

Since *Miscanthus*’ performance varies substantially across environments, understanding the genotype adaptability represents a significant challenge for breeders. Hence, it is crucial to understand the response patterns due to the G×E interaction to accelerate the selection process and to reduce the phenotyping costs associated with field evaluation experiments (Jarquin et al., 2014; Crossa et al., 2017). Breeding progress depends on identifying genotypes that outperform elite varieties in a target population of environments (TPE) or in a wide range of environmental conditions (Cooper et al., 2023).

Widener et al. (2024) used the same *M. sacchariflorus* and *M. sinensis* datasets that are the subjects of the current study to compare the predictive ability of two GS models; one included only main effects, while the other also included the G×E interaction. Both models were contrasted considering the most challenging prediction scenario which is mimicked by the cross-validation scheme CV00 (predicting untested lines in novel environments). These authors did not find significant improvements in predictive ability by including the G×E interaction compared to the main effects models. However, there are not cross-validation schemes developed for perennial crops and those developed for annual crops have been adapted.

In this study, within two *Miscanthus* species, we assess how the ability to predict biomass yield using different prediction models that account for various G×E interaction components perform under different cross-validation schemes. Cross-validation schemes were designed specifically for perennial crops considering as starting point the classic cross-validation schemes of: CV2, i.e. predicting tested genotypes in observed environments; CV1, i.e. untested genotypes in observed environments; CV0, i.e. tested genotypes in novel environments; and CV00, i.e. untested genotypes in novel environments (Persa et al., 2021). In addition, forward prediction was tested i.e. predicting the biomass yield for subsequent years using data from previous evaluation years. For all the previous cross-validation schemes, to be in accordance with the nature of the perennial crops that can be harvested more than once, we redefined the concept of environment to site-by-harvest year combination.

## 2 MATERIALS AND METHODS

### 2.1 Genomic Data

The *M. sacchariflorus* and *M. sinensis* panels studied here have been previously documented by Clark et al. (2014) and Clark et al. (2019c). Originally, for *M. sacchariflorus*, 722 clones were initially considered (Njuguna et al., 2023). This panel underwent genotyping and filtering to eliminate SNPs with more than 50% missing data and a minimum minor allele frequency of 0.01 (Głowacka et al., 2012). Applying quality control of the dataset resulted in a refined dataset comprising 34,605 SNP markers and 516 unique clonal genotypes available for subsequent analyses.

In the case of *M. sinensis*, the SNP markers were derived from RadSeq data, initially filtered for 568 genotype clones, and a sequence of quality control steps were applied. First, SNPs with a minimum cell rate of 0.04 and a minimum minor allele frequency of 0.02 were discarded. Second, individuals with over 30% missing data and minor allele frequencies below 0.05 were excluded (Widener et al., 2024). This resulted in a final set of 46,177 SNP markers and 260 unique *M. sinensis* clonal genotypes being available for analysis.

### 2.2 Phenotypic Data

A detailed description of the available phenotypic data sets is provided in Clark et al. (2019c), Njuguna et al. (2023) and Widener et al. (2024), with a brief overview being provided here.

Dataset 1 (*Msa*): Comprises a total of 7,740 phenotypic field records derived from 516 unique *M. sacchariflorus* clonal genotypes established in 2015 and evaluated in 2016, 2017 and 2018. These trials were grown in four temperate locations at higher altitudes – Sapporo, Japan by Hokkaido University (HU); Urbana, IL, USA by the University of Illinois (UI); Chuncheon, South Korea by Kangwon National University (KNU); Foulum, Denmark by Aarhus University (AU); and in one subtropical location at lower altitude – Zhuji, China by Zhejiang University (ZJU) (Njuguna et al., 2023; Widener et al., 2024). Biomass yield and 15 other yield trait components were recorded in all locations and years.

Dataset 2 (*Msi*): Comprises a total of 1,870 phenotypic field records derived from 260 unique *M. sinensis* clonal genotypes established in 2012 and evaluated in three temperate locations, namely, Sapporo, Japan by Hokkaido University (HU); Leamington, Canada by New Energy Farms (NEF); Chuncheon, South Korea by Kangwon National University (KNU); and one subtropical location Zhuji, China by Zhejiang University (ZJU) (Clark et al., 2019c; Widener et al., 2024). Biomass yield and 14 other yield trait components were evaluated.

A randomized complete block design with two to three replications at each site was used. For each trial, individual plants were grown and spaced two meters apart. Biomass yield was measured by collecting the stems 15-20 cm above the ground during the harvest season in November or December. Then, these were dried at 60°C until they reached a constant dry weight. The yield was expressed in Megagrams per hectare (Mg ha−1), and the dry weight of each plot was divided by the area of the plots (Njuguna et al., 2023; Clark et al., 2019c).

### 2.3 Statistical Models

Six different linear models were tested, which made predictions based on the site (S), harvest time (T), and genotype (G) effects, plus some combination of their interactions. A baseline, main effects-naive model that included line (L) rather than genomic information (M1: S+T+L), was compared against a baseline STGBLUP (M2: S+T+G) model, which also includes the main effect of the marker SNPs via covariance structures, and four models including different types of genotype-by-environment (G×E) interaction terms that are different to the conventional models evaluated in annual crops (M3: S+T+G+G×S; M4: S+T+G+T×S; M5: S+T+G+G×T; and M6: S+T+G+G×S+G×T+T×S+G×T×S) Details of these models and their corresponding effects are provided below.

#### 2.3.1 Baseline main effects naïve model

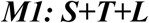

Consider that *y*_*ijk*_ represents the biomass yield value of the *i*^th^ (*L*_*i*_; *i* = 1,2,.., *n*) genotype observed in the *j*^th^ (*S*_*j*_; *j* = 1,…, *J*) site and measured in the *k*^th^ (*T_k_*; *k* = 1, 2, 3) time with 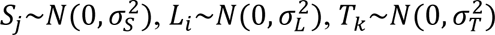, and 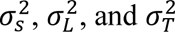 representing the corresponding variance components. In this model, the year effect (*T*_k_) was considered to allow borrowing the information within the sites but being harvested at different years and extra care was taken to avoid contamination of the training set (details provided below in the cross-validation section). Hence, the phenotypic trait is described as the sum of a constant common effect across the lines, sites, and harvest time (µ) plus random effects due to the site stimuli (*S_j_*), the harvest time (*T_k_*), and the corresponding genotype (*L_i_*), and a random effect (*ε_ijk_*) capturing the unexplained variability of the model with 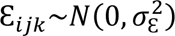. The resulting linear predictor is as follows.

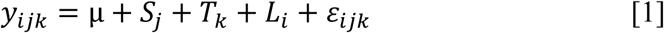

One disadvantage of this model is that the genotype (*L_i_*) effect relies only on phenotypic information and does not allow the borrowing of information between the observed and the non-observed genotypes. To overcome this limitation, the genomic information of the individuals is considered to connect phenotyped and un-phenotyped individuals (Persa et al., 2021)

#### 2.3.2 Baseline model including marker effect

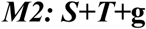

For including the genomic information, here, the main effects of the marker SNPs, *b*_*m*_ are modeled as independent and identically distributed (IID) random variables following a normal distribution centered at zero, all having a common variance such that 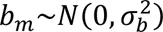 with 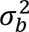 as the corresponding variance component. Hence, a linear combination between *p* markers, *x_im_* (*m* = 1,…, *p*) and their corresponding effects *b_m_* is used to represent the genomic effect of the *i*^th^ genotype as follows: 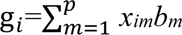. Here g represents the approximation of the true genetic value; *x*_*im*_ is the genotype value {0, 1, 2} of the *i*^th^ individual at the *m*^th^ marker. Following the results of the multivariate normal distribution (Persa et al., 2021), we have the vector of genomic effects 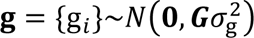, where ***G*** is the kinship matrix whose entries describe the genomic similarities between pairs of genotypes and 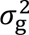 is the corresponding variance component. The other model terms remain as described before. Collecting the previous results, we have that the resulting linear predictor is as follows

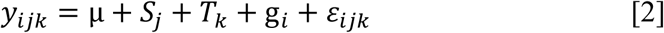

This model allows the borrowing of the information between the phenotype and the un-phenotype individuals, hence enabling the prediction of the unobserved individuals. On the other hand, a limitation of this model is that the genomic value g*_i_* of the *i*^th^ genotype is the same across the environment upon an environmental constant, such that the effect of the different environments is not captured.

To overcome this limitation, the reaction norm model was implemented allowing specific genomic effects at each set of environmental conditions.

#### 2.3.3 Incorporating the interaction between the markers (g) and site (S)

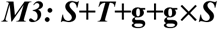

The models above accounted for the main effects of markers (g) and the main effects of the site without accounting for the interaction between these. In this model M3, the interaction between each marker SNPs and each site accounts for the genotype-by-site **g**×***S*** interaction. The vector of genotypes and sites interaction effects, ***gS*** = {*gS_ij_*}, is modeled via variance-covariances structures such that 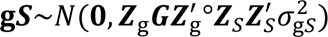, with a mean of zero and 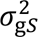 as the corresponding variance component (Jarquin et al., 2014). Where the ***Z***_g_ and ***Z****_S_* are the incidence matrices that connect phenotypic observations with genotype and sites, respectively, and “°” represents the cell-by-cell product between the two matrices, also known as the Hadamard product or the Shur product, which is an element-wise multiplication of two matrices of the same size (Styan, 1973). The resulting linear predictor is

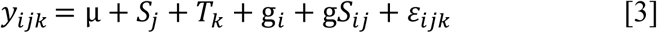

Here, the G×S interaction term allows the borrowing of information across the sites for specific genomic effects on each site.

#### 2.3.4 Main effect and harvest-by-site interaction model

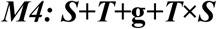

Here, the interaction component of G×S from the previous model was replaced by ***T***×***S***, which represents the interaction term of harvest time (***T***) and site (***S***), resulting in a vector containing sort of genotype-by-site multiplicative effects. The vector of interaction effects between time and harvest *TS* = {*TS*_*jk*_} is assumed to follow a normal distribution ***TS*** = 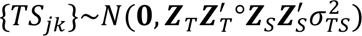, with a mean centered on zero and 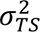 as the corresponding variance component and **Z***_T_* is the incidence matrices that connect the phenotype observations to harvest time, resulting in a sort of environment. The resulting model becomes

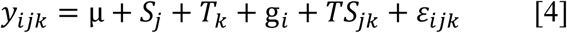

This model attempts to explicitly explain how the harvest time affects the biomass yield across sites.

#### 2.3.5 Main effect and genotype-by-harvest interaction models

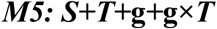

The marker SNPs and time of harvest interaction (**g**×***T***) were considered in this model via covariance structures. The vector of genomic effects in interaction with time of harvest (*gT_ik_*) is expressed as 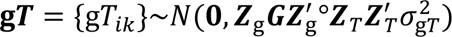, and it is assumed to follow a normal distribution with mean zero and 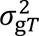 as the corresponding variance component. The resulting linear predictor is

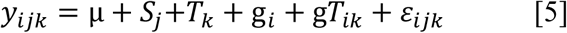

#### 2.3.6 A comprehensive model with all interactions

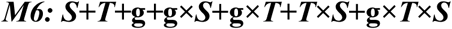

This is the most elaborate model, which includes all the main effects, and all the interactions as previously mentioned (**g**×***S*** genotype-by-site interaction; ***T***×***S*** harvest-by-site interaction; **g**×***T*** SNPs-by-harvest interaction). In addition, a three-way interaction (g*TS*_*ijk*_), which represents the interaction between the (**g**×***T***×***S*** SNPs-by-time of harvest-by-site interaction) was included. This model accounts for the complex genotype responses that can vary across different combinations of time points across different sites. Since the response of the same genotypes varies across the different years for the same site, the definition of environment is slightly distinct from the conventional site-by-year combination.

Our definition of environment is tied to the specific harvest at a given site (harvest 1, 2 or 3). Thus, a comparison can be made at the environmental level only between “environments” corresponding to the same harvest (e.g., harvest 1 for all sites, harvest 2 for all sites, and harvest 3 for all sites). Here, the three-way interaction is introduced using a covariance structure as follows

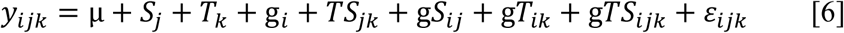

where, 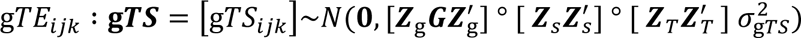 with mean **0** and 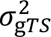 is the variance component and the other model terms as previously defined.

All the statistical analyses were done using the R software (R Core Team, 2021). The R package “BGLR” was used for fitting the described models (Pérez and de los Campos, 2014).

### 2.4 Cross-validation

To evaluate the levels of predictive ability that can be achieved implementing the described models, five different cross-validation scenarios were considered. These correspond to different prediction problems of interest for breeders. In all cases, the conventional cross-validation schemes implemented for annual crops were redesigned to be adapted to the needs for perennial species. Figure 1 introduces all the proposed cross-validation schemes. For an easy understanding, consider that all genotypes are observed in all sites across all years (harvests). The genotypes in orange/bright color represent the training set, those in gray color correspond to the testing set, and those in white color correspond to sets of genotypes that were not considered in the training nor the testing set.

**Figure 1.**
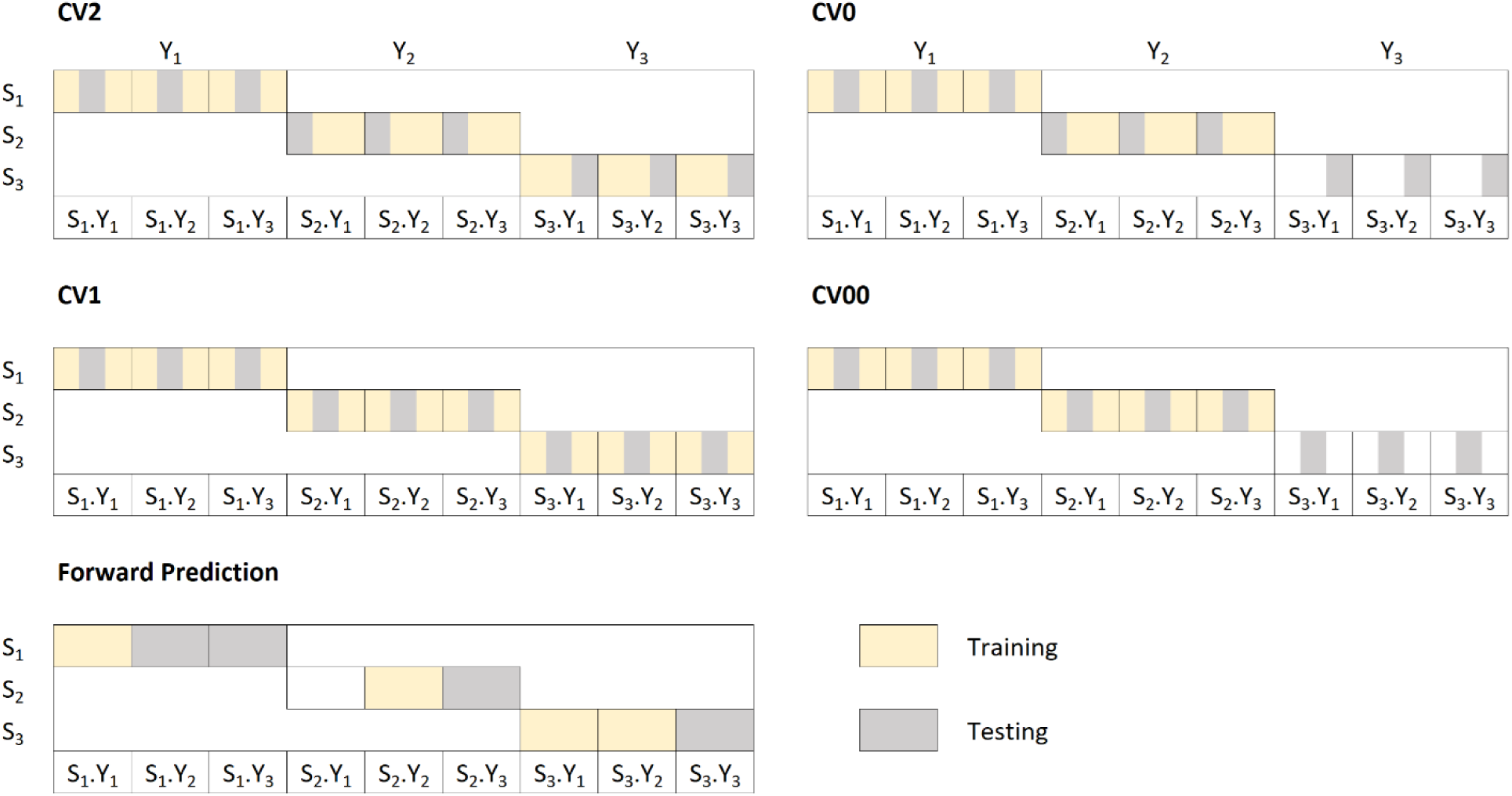
Graphical representation of five cross-validation schemes considered to predict trait performance of two *Miscanthus* populations (*Msa* and *Msi*) observed in multiple sites (S_1_, S_2_, and S_3_) and years (Y_1_, Y_2_, and Y_3_) with all genotypes observed in all sites, for a total of nine different combinations between sites and years (S_1_.Y_1_, S_1_.Y_2_, S_1_.Y_3_, S_2_.Y_1_, S_2_.Y_2_, S_2_.Y_3_, S_3_.Y_1_, S_3_.Y_2_, S_3_.Y_3_). CV2: Predicting tested genotypes in observed sites; CV1: Predicting untested genotypes in observed sites; CV0: Predicting tested genotypes in novel sites; CV00: Predicting untested genotypes in novel sites; Forward Prediction: Predicting the performance of tested genotypes in future years on the same site. The orange/bright color represents the training set, the gray color corresponds to the testing set, and the white color represents genotypes not considered either for training or testing sets.

The first four cross-validation scenarios attempt to split the data into training and testing sets, implementing a k-fold (k=5) approach. There, five-folds are created containing each of these around 20% of the data. The assignation depends on the implemented cross-validation scheme. For the validation, four folds (∼80% training) are considered for model training to predict the remaining fold (20% testing). Further details of these four cross-validation schemes are provided below.

#### 2.4.1 CV2: predicting tested genotypes in observed sites

CV2 (predicting tested genotypes in observed sites) represents incomplete field trials (Persa et al., 2021), and the target is the prediction of the genotypes that were evaluated in some sites but not in others (Figure 1). Here, since there is information from the same genotype for the three years of evaluation at each site, extra care was taken to assign the phenotypes of a genotype into the same partition or fold. Such that phenotypic records of those genotypes in the testing set can be only found in the training set but observed in other sites. This means that within the same site, across years, the three phenotypic records (year 1, year 2, and year 3) from the same genotype are assigned to the same fold. In this way, further contamination issues/inconsistencies of the training set are avoided. For example, predicting within the same site responses of genotypes in year 1 using phenotypic information from year 2 and/or year 3.

In this scheme, to conduct the assignment of phenotypic records to folds, we proceed as follows: first, we use the phenotypic information of year 1 only to assign phenotypes to folds; then, for the subsequent years, we use the same fold assignation. For example, if genotype 1 in site 2 was assigned to fold 2 (out of 5) in year 1, then the phenotypic information of this genotype in years 2 and 3 will be assigned to fold 2 for this site. The models are then implemented predicting one-fold at a time using the remaining 4 as a calibration set until all the folds are predicted. A total of 10 replicates were considered, such that the number of times the folds are predicted is 50 (5×10).

#### 2.4.2 CV1: predicting untested genotypes in observed sites

CV1 (predicting untested genotypes in observed sites) corresponds to predicting genotypes that have not been observed at any site (Figure 1). It is implemented for assessing the performance of untested genotypes in sites where other genotypes have been already observed. Here, the genotypes were randomly assigned to five folds. Similarly, in the previous case, each fold was used as a testing set, while the remaining four folds were used as a training set. This procedure was also repeated 10 times.

#### 2.4.3 CV0: predicting tested genotypes in unobserved sites

CV0 (predicting tested genotypes in unobserved sites), considers the prediction of the genotype performance in unobserved/novel sites (Figure 1). The phenotypic data from the same genotypes and/or other genotypes evaluated in different sites were utilized for model training. This prediction framework is built on top of the fold assignment of the CV2 scheme. Here, the phenotypic information of one site is masked as missing, but the target is to predict only one of the folds using the information from the other sites. We followed a similar scheme to the one proposed by Persa et al. (2021) to preserve comparable training set sizes across cross-validation schemes. This procedure was repeated for each fold at each site for all sites for 10 replicates generated in the CV2 scheme for a total of 250 (5×5×10) sub-fold-site partitions being predicted for each model.

#### 2.4.4 CV00: predicting untested genotypes in novel sites

CV00 (predicting untested genotypes in novel sites) is the most challenging prediction scheme, which mimics the real-life scenario of evaluating new cultivars in new sites (Figure 1). It combines the challenges of CV1 and CV0 scenarios involving the prediction of new clonal genotypes that have not been evaluated at any site yet in a site where no other genotypes have been observed either. This scenario is built up based on the fold assignment of the CV1 scheme, and similarly to CV0, we follow the same cross-validation design proposed by Persa et al. (2021). Here, one fold is predicted at a time after discarding the phenotypic information of the target site and the corresponding phenotypes of those genotypes in the current fold but observed in the training sites. This procedure was repeated for each fold at each site for all sites for 10 replicates generated in the CV1 scheme for a total of 250 (5×5×10) sub-fold-site partitions being predicted for each model.

#### 2.4.5 Forward prediction: predicting future harvests

Forward Prediction (predicting future harvests), here, the already observed data at early stages of the field testing was used to predict the performance of the subsequent breeding years in an attempt to reduce the phenotyping costs and, more importantly, the length of the selection cycle (Figure 1). This prediction scheme holds the potential to evaluate genotypes at early stages for selecting candidate breeding parents to be crossed within and between the studied species.

Here, genomic prediction models were implemented for evaluating three different situations: *i*) model calibration using phenotypic data of the first year to predict the second and third years; *ii*) model calibration using phenotypic data from the second year for predicting the third year; and *iii*) model calibration using the pooled data of the first and the second year to predict the performance at the third year of evaluation for those sites where data was available for the three years.

### 2.5 Assessment of Prediction Accuracy

As mentioned before, our definition of environment is a little different from the conventional considered for annual crops. This change was necessary because the number of years the genotypes have been planted in fields should be also considered to allow a fair comparison between the different stages (years). In our case, the environment is defined as the site-by-harvest year combination. For this reason, the model’s performance was evaluated by calculating Pearson’s correlation between the predicted and the observed values within each site-by-harvest year combination, resulting in an *environment*. The weighted average correlation across the environments was estimated to account for the uncertainty and sample size differences for the different cross-validation schemes according to Tiezzi et al. (2017)

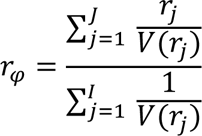

where *r*_*j*_ represents Pearson’s correlation between the observed and predicted values at the *j^th^ environment* ( *j*=1, 2,…, *J*; *J* = 15 for *Msa* and *J* = 10 for *Msi*), with 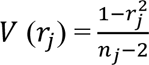 representing the sampling variance and *n*_*j*_ is the corresponding number of observations.

### 2.6 Variance components

Variance components are crucial to assess the contribution of the different main (genotype, site, and time) and interaction effects for explaining trait variability. For each model term, the percentage of explained variability was computed. Here, the proportion of variability addressed by the *o*^th^ {*o=*1,2,…,*O*} model term is calculated by dividing its corresponding specific variance by the sum of all variance components in the model multiplied by 100.

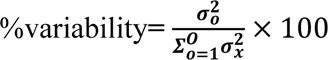

## 3 RESULTS

### 3.1 Dataset 1 - Miscanthus sacchariflorus (Msa)

#### 3.1.1 Variance components

The computation of the variance components of the different model terms is conducted by considering a full data analysis (i.e., no phenotypic records are masked as missing values). Table 1 presents the percentage of variability explained by each model term for all models for the *Miscanthus sacchariflorus* population. For the main effects model M1 (***S***+***T***+***L***), which is based solely on the phenotypic information, the site main effect (S) explains 15.6% of the total variability, while the line main effect (L) captures 23.4%, and the time main effect only 3.6%, the residual term (R) addresses 57.5% (unexplained variability). When the main effect of the markers is introduced in M2 (***S***+***T***+**g**) via the covariance structure ***G***, it captures 8.1% of the variability. Meanwhile, the site (***S***) and residual (**R**) variances increase to 18.9% and 68.3%, respectively, compared with M1, while the variance component of T remains practically unchanged (∼4.7%). The interaction between the markers and sites (**g**×***S***) included in M3 captures 14.6% of the total variability and reduces the residual variance (**R**) to 52.4%. The interaction between time of harvest and site (***T***×***S***) in M4 captures 35.8% of the total variability, and the residual term was increased to 56.2%.

**Table 1.** Percentage of explained variability *σ*^2^ by each model term for six models (M1 *S*+*T*+*L*; M2 *S*+*T*+g; M3 *S*+*T*+g+g×*S*; M4 *S*+*T*+g+*T*×*S*; M5 *S*+*T*+g+g×*T*; M6 *S*+*T*+g+g×*S*+*T*×*S*+g×*T*+g×*T*×*S*) of *Miscanthus sacchariflorus* collection comprised of 516 unique genotypes observed in 5 sites and evaluated for three years. The model terms of the six considered models are as follows: *S*, main effect of sites; *L*, main effect of genotypes; g, main effect of markers; *T*, main effect of time; g×*S* corresponds to the interaction between marker SNPs and sites; *T*×*S* for the interaction between the time of harvest and the sites; g×*T* for the interaction between the marker SNPs and time of harvest; and g×*T*×*S* for the three-way interaction between the marker SNPs, time of harvest, and site.

| Model | $\hat{\sigma}_S^2$ | $\hat{\sigma}_T^2$ | $\hat{\sigma}_L^2$ | $\hat{\sigma}_g^2$ | $\hat{\sigma}_{T \times S}^2$ | $\hat{\sigma}_{g \times S}^2$ | $\hat{\sigma}_{g \times T}^2$ | $\hat{\sigma}_{g \times T \times S}^2$ | $\hat{\sigma}^2$ |
| --- | --- | --- | --- | --- | --- | --- | --- | --- | --- |
| M1: $S+T+L$ | 15.6 | 3.6 | 23.4 | | | | | | 57.5 |
| M2: $S+T+g$ | 19.0 | 4.7 | | 8.1 | | | | | 68.3 |
| M3: $S+T+g+g \times S$ | 22.8 | 4.7 | | 5.6 | | 14.6 | | | 52.4 |
| M4: $S+T+g+T \times S$ | 0.0 | 0.0 | | 7.9 | 35.8 | | | | 56.2 |
| M5: $S+T+g+g \times T$ | 18.1 | 4.5 | | 7.8 | | | 1.1 | | 68.5 |
| M6: $S+T+g+T \times S+g \times S+g \times T+g \times T \times S$ | 0.0 | 0.0 | | 4.2 | 40.5 | 20.7 | 1.0 | 10.5 | 23.1 |

Interestingly, when this interaction was introduced in models M4 and M6, the variability captured by the site and time of harvest terms was reduced to practically zero, highlighting its importance. Hence, the phenotypic data across the years shows heterogeneity of variances. The average phenotypic variability of the sites in year 1 was 13.3, which is the lowest. The second year exhibited a phenotypic variability of 14.5, whereas the third year showed the highest variability, 29.35.

In model M5, the interaction between marker SNPs and harvest time was included, and it captured 1.1% of the phenotypic variability. The corresponding values of the other terms remain within the range of values shown in models M1-M3, except for the residual term that addressed the largest variability among all models (∼68.5%).

Furthermore, the most elaborate model M6, which, besides including all of the previous main and interaction effects, also considers the three-way interaction **(g**×***T***×***S***) between marker SNPs-site-harvest time, reduced the residual variance the most to only 23.1%. Here, similarly to model M4, the variance component of site and time effects was reduced to practically zero while the harvest time-by-site interaction captured ∼40.5% of the variability and the three-way interaction ∼10.5%.

#### 3.1.2 Assessment of prediction accuracy

As mentioned before, the model’s ability to deliver accurate predictions is assessed on a trial basis as the Pearson correlation between predicted and observed values within the same site-harvest time combination across different cross-validations scenarios (CV2, CV1, CV0, CV00). The results for *M. sacchariflorus* are shown in Supplementary Table S1 (A, B, C and D). Table 2 presents, for 10 replicates, the average mean correlation (corresponding standard deviation values) and average mean square error values for four cross-validation schemes and six prediction models (M1 ***S***+***T***+***L***; M2 ***S***+***T***+**g**; M3 ***S***+***T***+**g**+**g**×***S***; M4 ***S***+***T***+**g**+***T***×***S***; M5 ***S***+***T***+**g**+**g**×***T***; M6 ***E***+***T***+**g**+***T***×***S***+**g**×***S***+**g**×***T***+**g**×***T***×***S***) implemented for predicting biomass yield.

**Table 2.**
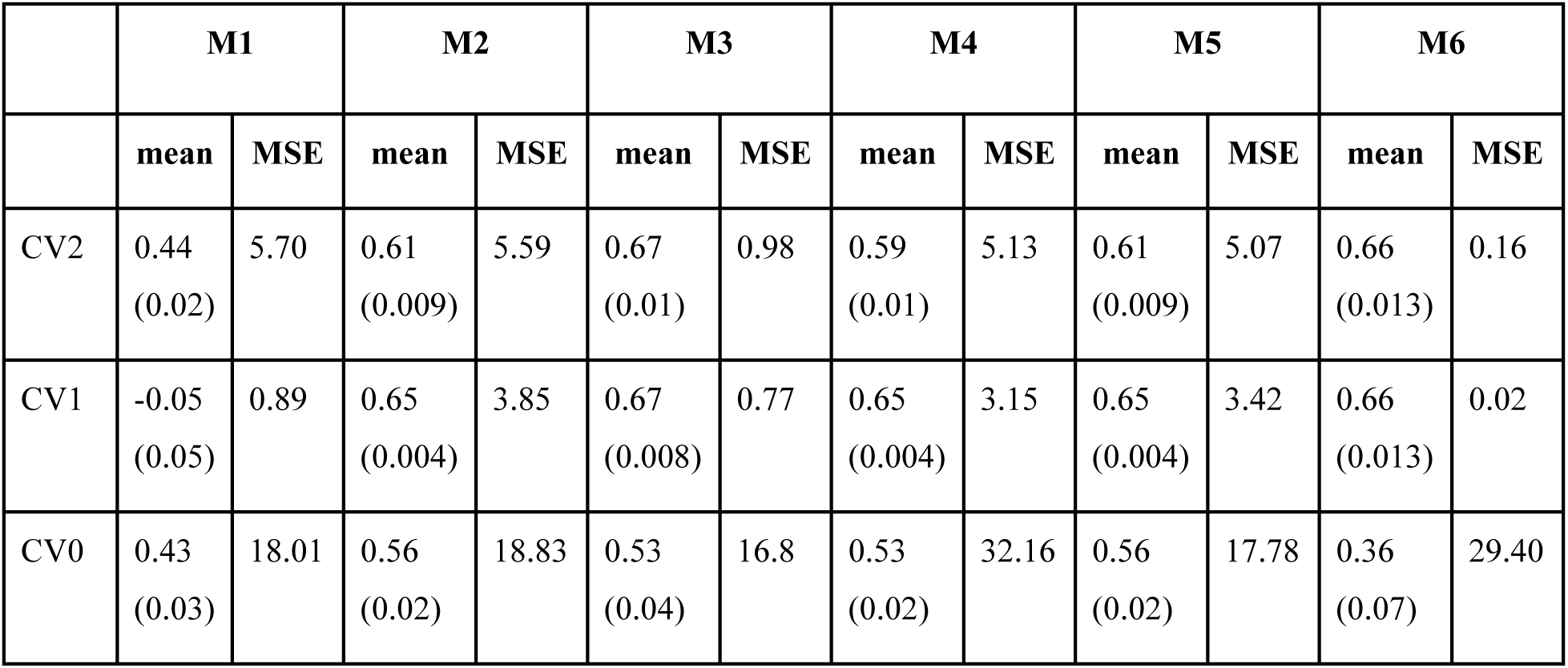
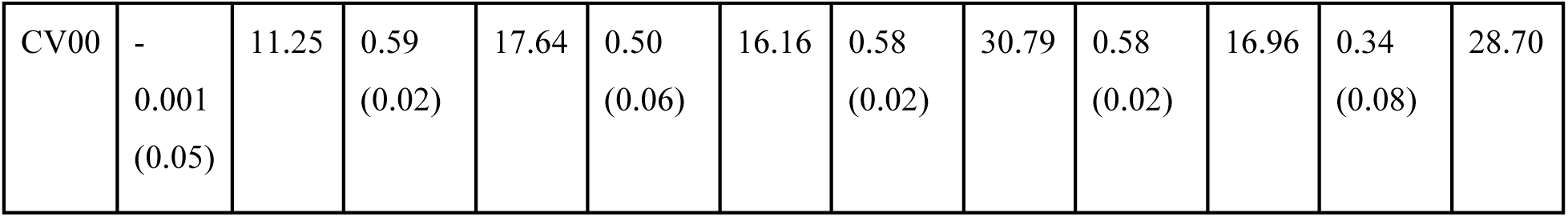
Average mean correlation (corresponding standard deviation) and average mean square error values for six different genomic prediction models (M1-M6) implemented on a *Miscanthus sacchariflorus* population for predicting yield performance under four different cross-validation schemes (CV2, CV1, CV0, and CV00). The results correspond to 10 replicates of five-fold cross-validation. M1: *S*+*T*+*L*; M2: *S*+*T*+*g*; M3: *S*+*T*+**g**+**g**×*S*; M4: *S*+*T*+g+*T*×*S*; M5: *S*+*T*+g+g×*T*; and M6: *S*+*T*+g+*T*×*S*+g×*S*+g×*T*+g×*T*×*S*. *S*, *T*, *L* and g correspond to the main effect of the site, harvest time, genotype effect, and marker SNPs; g ×*S*, *T*×*S*, g×*T*, and g×*T*×*S* correspond to the interaction effects between marker SNPs and site, harvest time and site, marker SNPs and harvest time, and the three-way interaction between marker SNPs and harvest time and site.

In CV2, the baseline model M1 showed the lowest average mean correlation (∼0.44, SD=0.02) and the highest MSE (∼5.7). When the marker SNP information was considered in M2, the correlation increased to ∼0.61 (SD=0.01) for an MSE of 5.59. This represents a relative improvement in the predictive ability of around 39% with respect to M1, while the MSE values remained almost the same. The model that includes the interaction between markers and sites in M3 improves the predictive ability by around 10% with respect to the baseline genomic prediction model M2. However, a more significant improvement was accomplished in terms of the MSE, which was reduced by around 90% compared to M2. The models, M4 and M5, that integrate interactions between harvest time and site and between marker SNPs and sites, respectively, returned similar average mean correlations of around ∼0.6 (SD=0.01) and an average MSE of ∼5.1. The most complex model, M6 presented an average mean correlation of ∼0.66 (SD=0.013) and the lowest average MSE across models (∼0.16). This represents a significant reduction in error of more than 97% (0.16/5.59) with respect to model M2 and of around 84% (0.16/0.98) with respect to M3. Hence, model M3, which included the **g**×***S*** interaction, expressed the highest mean average correlation of 0.67 with an MSE of 0.98, whereas model M6 showed a similar prediction accuracy of 0.66 with a significantly lower MSE value of 0.16.

In CV1, as expected, the model M1, which does not include genomic information, showed an average correlation close to zero (−0.05). When the main effects of the marker SNPs were included in model M2, the predictive ability was 0.65 (SD=0.004) for an MSE of 3.85. Interestingly, contrary to expectations, the predictive ability and MSE of the models for CV1 were better (higher correlations and reduced bias) than CV2, except for M1. Usually, opposite trends are expected when the genotypes are already in some environments. When the different G×E interactions **(g**×***S***, ***T***×***S***, **g**×***T***, **g**×***T***×***S***) were included in the models, the correlations increased, ranging from 0.65 to 0.67 for models M3-M6. The models M3 (0.67) and M6 (0.66) showed slightly better predictive ability values than the other models (M4 and M5, 0.65); however, the average MSE was significantly reduced with M6 (0.02) compared to M3 (0.77). Although the baseline M2 model performed similarly in terms of predictive ability compared to M6, its MSE was significantly larger (3.85 vs. 0.02).

Under the CV0 scenario, M1 returned an average mean correlation of 0.43 (SD=0.03) and an MSE of 18.01. Here, M2 and M5 showed the highest average mean correlation (∼0.56), with M5 returning a slightly reduced MSE value (17.78 vs. 18.83). In general, M3 returned the lowest MSE across all models (16.8).

For the most complex prediction scenario CV00, again and as expected, M1 returned an average correlation close to zero. The model that includes the main effect of the markers M2 (0.59) slightly outperformed the other models M3-M6 that accounts for interactions varying between 0.34 to 0.58. Regarding the average MSE, M1 returned the lowest value (11.25); however, due to its null predictive power, this model was discarded. Of the remaining models showing moderate to high correlations, the MSE varied between 16.16 (M3) and 30.79 (M4). In this scenario, the most complex models based on both criteria (high correlation and reduced MSE) are M2 and M5, with M3 being more parsimonious (fewer terms).

Across cross-validation scenarios, the predictive ability in CV2 and CV1 benefited from the inclusion of interaction terms. Whereas in CV0 and CV00, the main effects model, including marker SNPs only (i.e., M2), performed better than the models including interactions.

#### 3.1.3 Assessment of forward prediction

The forward prediction scenario allows the assessment of the model’s predictive ability of genotype performance in advanced growing/developmental stages using phenotypic records from observed at early stages (year 1, or years 1 and 2) for model calibration. As mentioned before, *Miscanthus* is a perennial species that typically requires phenotypic information from three years to generate reliable results. Hence, the accurate identification of superior cultivars at early stages is crucial for shortening the selection cycle. Here, the above-described six prediction models were considered for three different prediction situations.

Table 3 presents the mean predictive ability and MSE values across 15 site-by-harvest year combinations obtained for the six described models and three prediction scenarios. In the forward prediction scenario 1, the observed data of the genotypes from year 1 were used as a training set to predict the biomass yield of year 2 and year 3. Predicting data from year 2, M3 returned the highest correlation (0.666) among all models and the lowest MSE (11.31). Across models the correlation values varied between 0.581 (M1) and 0.666 (M2) and the MSE values between 11.31 (M3) to 16.51 (M6). Here, it is important to mention that the model that did not include genomic information, M1, returned a moderate to high correlation (0.581), highlighting the importance of the availability of phenotypic information collected at early stages. Regarding the prediction for year 3, the correlations were slightly reduced, and the MSE values increased around threefold, with model M2 returning the best results (0.573 and MSE=31.0). This last case is probably the most useful for breeders because it enables the identification of superior cultivars using only data from year 1. If the predictive ability of the models is high, it could help to shorten the selection process by two years.

**Table 3.** Mean predictive ability based on the Pearson correlation between predicted and observed values within site-by-harvest year combinations (15) and the corresponding mean square error for six different genomic prediction models in *Miscanthus sacchariflorus* for three different prediction scenarios. Models: M1 *S*+*T*+*L*; M2 *S*+*T*+*g*; M3 *S*+*T*+**g**+**g**×*S*; M4 *S*+*T*+g+*T*×*S*; M5 *S*+*T*+g+g×*T*; M6 *S*+*T*+g+*T*×*S*+g×*S*+g×*T*+g×*T*×*S* under forward prediction. *S*, *T*, *L* and g correspond to the main effect of the site, harvest time, genotype effect, and marker SNPs; g ×*S*, *T*×*S*, g ×*T*, and g ×*T*×*S* correspond to the interaction effects between marker SNPs and site, harvest time and site, marker SNPs and harvest time, and the three-way interaction between marker SNPs and harvest time and site. Prediction situations: i) TRN Y_1_ - TST Y_2_ & Y_3_ corresponds to the cross-validation situation of using year 1 as a training set for predicting year 2 and 3; ii) TRN Y_2_ - TST Y_3_ it describes the prediction situation of using year 2 as training set for predicting for year 3; iii) TRN Y_1_ & Y_2_ - TST Y_3_ represents the situation of utilizing the pooled data of year 1 and 2 as training set to predict the year 3.

| TRN $Y_1$ - TST $Y_2$ & $Y_3$ | | | | | | | | | | | | |
| --- | --- | --- | --- | --- | --- | --- | --- | --- | --- | --- | --- | --- |
| Mea<br>n | M1<br>($S+T+L$) | | M2<br>($S+T+g$) | | M3<br>($S+T+g+g \times S$) | | M4<br>($S+T+g+T \times S$) | | M5<br>($S+T+g+g \times T$) | | M6<br>($S+T+g+g \times S+T \times S+g \times T+g \times T \times S$) | |
|  | PA | MS<br>E | PA | MS<br>E | PA | MS<br>E | PA | MS<br>E | PA | MS<br>E | PA | MSE |
| $Y_2$ | 0.581 | 12.60 | 0.595 | 11.84 | 0.666 | 11.31 | 0.595 | 11.34 | 0.594 | 14.70 | 0.641 | 16.51 |
| $Y_3$ | 0.509 | 34.61 | 0.549 | 32.52 | 0.573 | 31.00 | 0.549 | 34.70 | 0.553 | 37.16 | 0.545 | 44.31 |
| TRN $Y_2$ - TST $Y_3$ | | | | | | | | | | | | |
| Y<sub>3</sub> | 0.56<br>7 | 23.5<br>5 | 0.58<br>2 | 21.7<br>6 | 0.65<br>4 | 16.2<br>5 | 0.58<br>2 | 29.2<br>8 | 0.58<br>2 | 26.4<br>7 | 0.634 | 24.83 |
| TRN Y<sub>1</sub> & Y<sub>2</sub> - TST Y<sub>3</sub> |  |  |  |  |  |  |  |  |  |  |  |  |
| Y<sub>3</sub> | 0.56<br>7 | 26.4<br>1 | 0.57<br>7 | 25.6<br>3 | 0.62<br>8 | 23.1<br>5 | 0.57<br>7 | 35.1<br>6 | 0.57<br>6 | 26.2<br>7 | 0.656 | 30.53 |

The forward prediction scenario 2 uses data from year 2 to predict year 3 only. Here, the model M1 based only on phenotypic information returned a correlation of 0.567 for an MSE of 23.55. Again, the most accurate model was M3 for a correlation value of 0.654 and an MSE of 16.25. The other models exhibited correlations ranging between 0.582 and 0.634 and MSE values between 21.76 and 29.28.

Combining data from years 1 and 2 to predict the performance of genotypes in year 3 under the forward prediction scenario 3 the model M1 based only on phenotypic information returned a correlation of 0.567 and a MSE of 26.41. The highest correlation (0.656) was obtained with M6 for an MSE of 30.53, while the lowest MSE (23.15) was produced with M3 model.

### 3.2 Dataset 2 - Miscanthus sinensis (Msi)

#### 3.2.1 Variance components

Similar patterns than for the *M. sacchariflorus* population were found for this population.

Especially, the variance components trends for the different models, and also the average phenotypic variability keeps similar ratios among years. Table 4 presents the variability percentage explained by each model term for all the models for the *M. sinensis* population. For model M1 (***S***+***T+L***), which depends on the phenotypic information only, the main effects of the site (***S***), time (***T***), and line (***L***) respectively captured 17.1%, 32.9%, and 9.3% of total variability. The inclusion of the main effects of marker SNPs (**g**) in model M2 captured 8.1% of the variability. In both models the unexplained residual variance remains at about 40% out of the total. Also, the variance component of the other terms, such as site and time, remains unchanged. The interaction between the markers and the sites **(g**×***S***) included in model M3, captures 13.0% of the total variability and gradually reduces the residual variance (**R**) to 31.1%. The interaction between the time of harvest and site (***T***×***S***) in model M4 captures about 52% of the total variability, while the residual term was increased to 37.8%. Meanwhile, the variance associated with the site (***S***) and time (***T***) terms was reduced to zero. Similarly to the previous population, the average phenotypic variance for each year across sites was the lowest for year 1 (0.64), while for the second and third year these values were 1.26 and 2.86, respectively. This represents a four-fold increase in year 3 compared to year 1. The interaction between the marker SNPs and time (**g**×***T***) is included in model M5 and captures only about 3% of the total variability. The corresponding values of the other terms remain within the range of values shown in models M1-M3. Furthermore, the most elaborated model, M6, which not only includes all the previous main and interaction effects but also considers the three-way interaction (**g**×***T***×***S***) between marker SNPs-harvest time-site, reduced the residual variance the most, to 14.0%. Similar to M4, the variance component of site and time effects was reduced to practically zero, while the harvest time-by-site interaction captured ∼51.6%, and the three-way interaction ∼6.2% of the total variability.

**Table 4.** Percentage of explained variability *σ*^2^ by each model term for six models (M1 *S*+*T*+*L*; M2 *S*+*T*+g; M3 *S*+*T*+g+g×*S*; M4 *S*+*T*+g+*T*×*S*; M5 *S*+*T*+g+g×*T*; M6 *S*+*T*+g+g×*S*+*T*×*S*+g×*T*+g×*T*×*S*) of *Miscanthus sinensis* collection comprised of 260 unique genotypes observed at 4 sites for evaluated for two to three years. The model terms of the six considered models are as follows: *S*, main effect of sites; *L*, main effect of genotypes; g, main effect of markers; *T*, main effect of time; g×S corresponds to the interaction between marker SNPs and sites; *T*×*S* for the interaction between the time of harvest and the sites; g×*T* for the interaction between the marker SNPs and time of harvest; and g×*T*×*S* for the three-way interaction between the marker SNPs, time of harvest, and site.

| Models | $\hat{\sigma}_S^2$ | $\hat{\sigma}_T^2$ | $\hat{\sigma}_L^2$ | $\hat{\sigma}_g^2$ | $\hat{\sigma}_{T \times S}^2$ | $\hat{\sigma}_{g \times S}^2$ | $\hat{\sigma}_{g \times T}^2$ | $\hat{\sigma}_{g \times T \times S}^2$ | $\hat{\sigma}^2$ |
| --- | --- | --- | --- | --- | --- | --- | --- | --- | --- |
| M1: $S+T+L$ | 17.2 | 33.0 | 9.4 | | | | | | 40.5 |
| M2: $S+T+g$ | 17.3 | 34.3 | | 8.1 | | | | | 40.2 |
| M3: $S+T+g+g \times S$ | 17.7 | 33.9 | | 4.4 | | 13.0 | | | 31.1 |
| M4: $S+T+g+T \times S$ | 0.1 | 0.1 | | 10.2 | 51.9 | | | | 37.8 |
| M5: $S+T+g+g \times T$ | 16.6 | 37.4 | | 7.2 | | | 2.9 | | 36.0 |
| M6:<br>$S+T+g+T \times S+g \times S+g \times T+g \times T \times S$ | 0.0 | 0.0 | | 3.4 | 51.6 | 21.3 | 3.5 | 6.2 | 14.0 |

#### 3.2.2 Assessment of prediction accuracy

The prediction accuracy for *M. sinensis* across different fivefold cross-validations schemes (CV2, CV1, CV0, CV00) are shown in Supplementary Table S2 (A, B, C and D). Table 5 presents for the 10 replicates, the mean average correlation, the corresponding standard deviations, and the average mean square error values for the four fivefold cross-validation schemes and the six aforementioned prediction models.

**Table 5.**
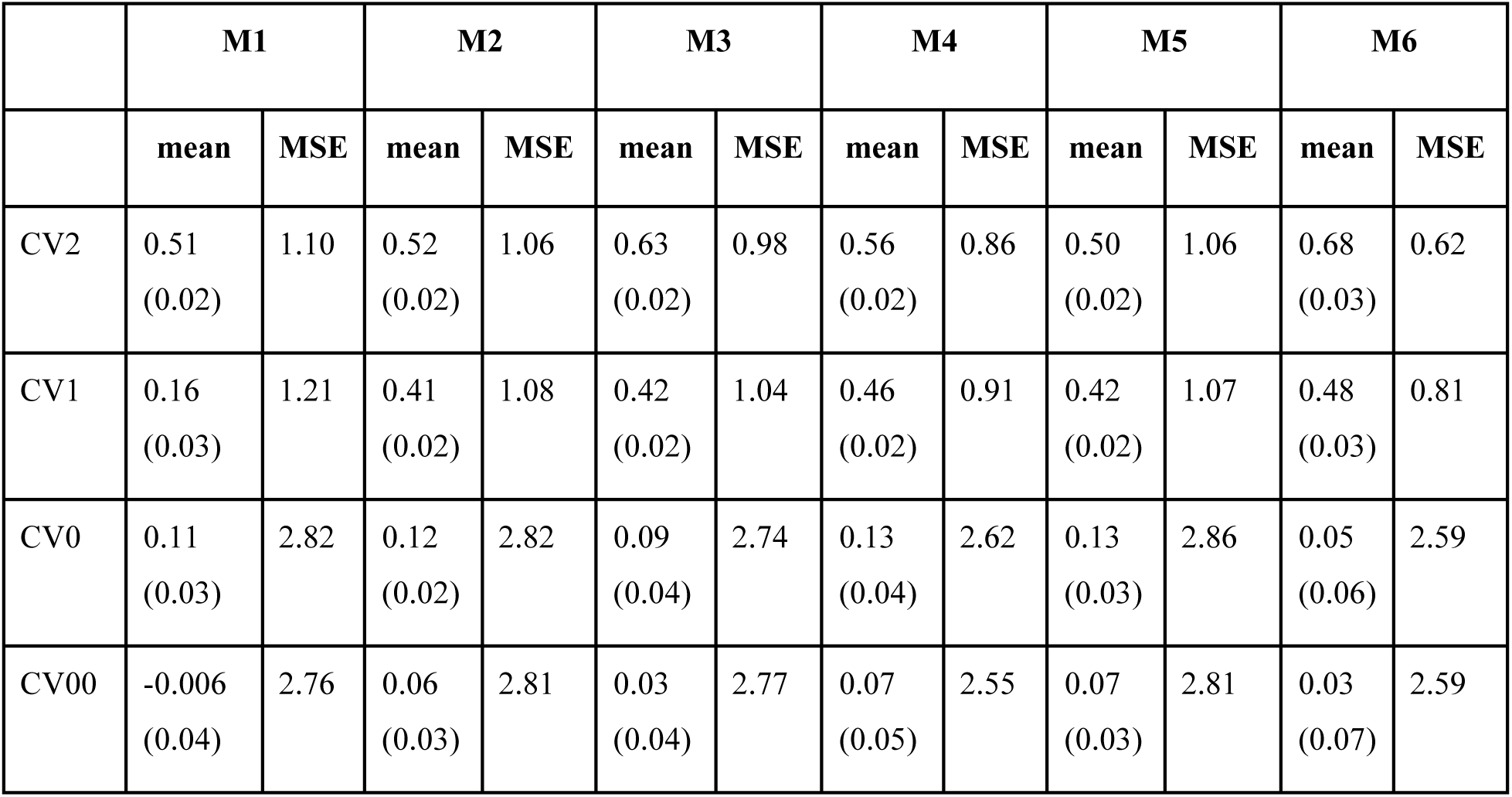
Average mean correlation (corresponding standard deviation) and average mean square error values for six different genomic prediction models (M1-M6) implemented on a *Miscanthus sinensis* population for predicting yield performance under four different cross-validation schemes (CV2, CV1, CV0, and CV00). The results correspond to 10 replicates of five-fold cross-validation. M1: *S*+*T*+*L*; M2: *S*+*T*+g; M3: *S*+*T*+**g**+**g**×*S*; M4: *S*+*T*+g+*T*×*S*; M5: *S*+*T*+g+g×*T*; and M6: *S*+*T*+g+*T*×*S*+g×*S*+g×*T*+g×*T*×*S*. *S*, *T*, *L* and g correspond to the main effect of the site, harvest time, genotype effect, and marker SNPs; g ×*S*, *T*×*S*, g×*T*, and g×*T*×*S* correspond to the interaction effects between marker SNPs and site, harvest time and site, marker SNPs and harvest time, and the three-way interaction between marker SNPs and harvest time and site.

|  | M1 |  | M2 |  | M3 |  | M4 |  | M5 |  | M6 |  |
| --- | --- | --- | --- | --- | --- | --- | --- | --- | --- | --- | --- | --- |
|  | mean | MSE | mean | MSE | mean | MSE | mean | MSE | mean | MSE | mean | MSE |
| CV2 | 0.51<br>(0.02) | 1.10 | 0.52<br>(0.02) | 1.06 | 0.63<br>(0.02) | 0.98 | 0.56<br>(0.02) | 0.86 | 0.50<br>(0.02) | 1.06 | 0.68<br>(0.03) | 0.62 |
| CV1 | 0.16<br>(0.03) | 1.21 | 0.41<br>(0.02) | 1.08 | 0.42<br>(0.02) | 1.04 | 0.46<br>(0.02) | 0.91 | 0.42<br>(0.02) | 1.07 | 0.48<br>(0.03) | 0.81 |
| CV0 | 0.11<br>(0.03) | 2.82 | 0.12<br>(0.02) | 2.82 | 0.09<br>(0.04) | 2.74 | 0.13<br>(0.04) | 2.62 | 0.13<br>(0.03) | 2.86 | 0.05<br>(0.06) | 2.59 |
| CV00 | -0.006<br>(0.04) | 2.76 | 0.06<br>(0.03) | 2.81 | 0.03<br>(0.04) | 2.77 | 0.07<br>(0.05) | 2.55 | 0.07<br>(0.03) | 2.81 | 0.03<br>(0.07) | 2.59 |

In the CV2 scheme, the baseline model M1 showed an average mean correlation of 0.51 (SD=0.02) and the highest MSE value (1.10). After including the SNPs in model M2, no significant improvements were observed returning an average mean correlation of 0.52 (SD=0.02) and an MSE value of 1.06. When the interaction between markers and sites is included in model M3, it improves the predictive ability by 21% (0.63/0.52×100-100) while its corresponding MSE was reduced by nearly 8% when compared to the basic genomic prediction model M2. Models M4 and M5, which integrate the interactions between harvest time and site and between marker SNPs and sites, showed average correlations ranging from 0.50 to 0.56, with the same standard deviation (SD=0.02). Compared to model M3, the MSE value of model M4 was reduced by 13% (0.86/0.98), while for model M5, it increased by 8% (1.06/0.98). The most elaborate model, M6 returned an average mean correlation of 0.68 (SD=0.03) and also showed the lowest MSE (0.62) value among all models. This represents a significant reduction of the MSE value of 42% (0.62/1.06) with respect to model M2 and around 37% (0.62/0.98) with respect to model M3. Hence, the most elaborated model, M6, expressed the highest mean average correlation of (0.68) and the lowest MSE value (0.62).

For the CV1 scheme, as expected M1 showed the lowest average correlation of 0.16 (SD=0.16) and the highest MSE value (1.21) among all the models. In theory, the average mean correlation of this model should be close to zero. The main effects of marker SNPs included in model M2 improved the predictive ability to 0.41 (SD=0.02) for a MSE value of 1.08. When the different interactions were included in the models, the correlations increased, ranging from 0.41 to 0.48 for models M3 to M6. Model M6 (0.48) showed the best predictive ability and the lowest MSE value (0.81).

Under the CV0 scheme, model M1 showed an average mean correlation of 0.11 (SD=0.03) and a MSE of 2.82. Here, models M4 and M5 returned the highest average mean correlation (0.13), with M4 having a slightly reduced MSE value of 2.62. Similar to the previous cross-validation schemes, model M6 returned the lowest MSE value (2.59) across all the models.

For CV00, model M1 returned an average correlation close to zero. The M4 and M5 models, which integrate interactions between harvest time and site and between marker SNPs and sites, showed the highest average correlation of 0.07, with M4 having a slightly reduced MSE value of 2.55. These models slightly outperformed M2, M3, and M6, whose correlations vary between 0.03 and 0.06 and their MSE values range from 2.59 to 2.81.

Across all four fivefold cross-validation schemes (CV2, CV1, CV0, and CV00) above mentioned, the predictive ability was benefited from the inclusion of interaction terms in some cases but not in others. The results from CV0 and CV00 showed the complications of predicting trait performance in novel environments.

#### 3.2.3 Assessment of forward prediction

Table 6 presents the results of the forward prediction scenario 1 where the observed data from the first year is used for predicting the biomass yield of the second year and the third year. Predicting data for year 2, model M1, which does not include the genomic information, showed the lowest predictive ability (0.41) and high MSE value (2.34). Whereas M6 returned the highest correlation (0.669) among all models with a reduced MSE (2.30). In addition, M3 showed a similar prediction accuracy (0.659) and the lowest MSE (1.90) among all the models thus being considered the best prediction strategy.

**Table 6.**
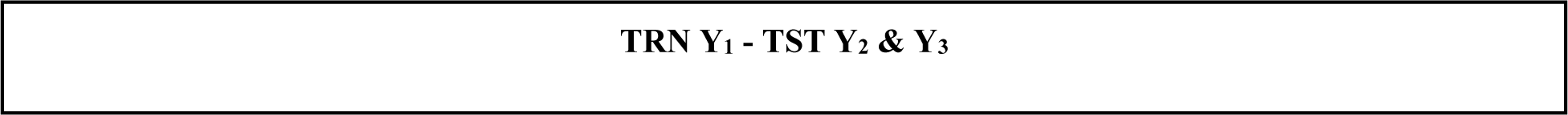
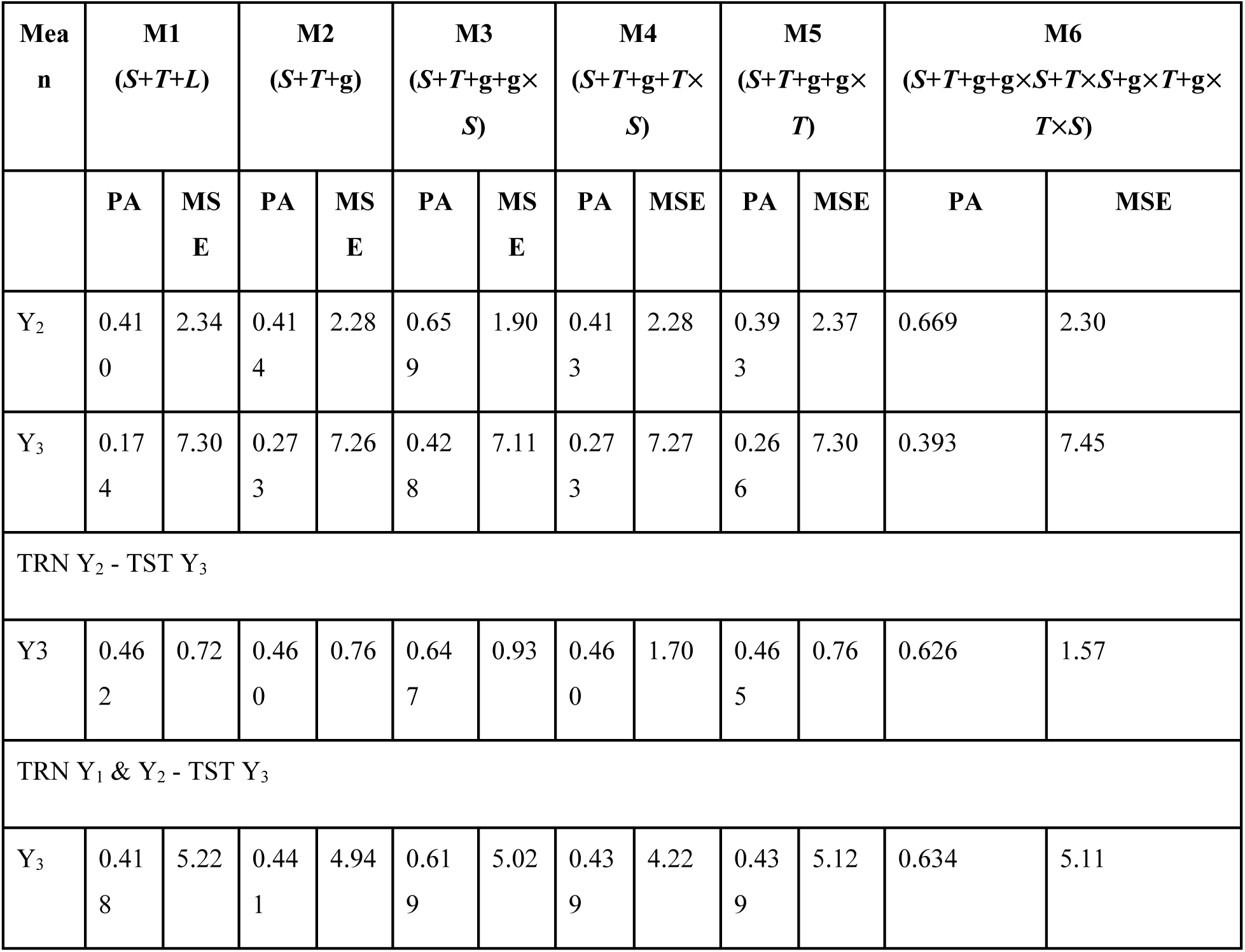
Mean predictive ability based on the Pearson correlation between predicted and observed values within site-by-harvest year combinations (10) and the corresponding mean square error for six different genomic prediction models in *Miscanthus sinensis* for three different prediction situations. Models: M1 *S*+*T*+*L*; M2 *S*+*T*+g; M3 *S*+*T*+**g**+**g**×*S*; M4 *S*+*T*+g+*T*×*S*; M5 *S*+*T*+g+g×*T*; M6 *S*+*T*+g+*T*×*S*+g×*S*+g×*T*+g×*T*×*S* under forward prediction. *S*, *T*, *L* and g correspond to the main effect of the site, harvest time, genotype effect, and marker SNPs; g ×*S*, *T*×*S*, g ×*T*, and g ×*T*×*S* correspond to the interaction effects between marker SNPs and site, harvest time and site, marker SNPs and harvest time, and the three-way interaction between marker SNPs and harvest time and site. Prediction situations: i) TRN Y_1_ - TST Y_2_ & Y_3_ corresponds to the cross-validation situation of using year 1 as a training set for predicting year 2 and 3; ii) TRN Y_2_ - TST Y_3_ it describes the prediction situation of using year 2 as training set for predicting for year 3; iii) TRN Y_1_ & Y_2_ - TST Y_3_ represents the situation of utilizing the pooled data of year 1 and 2 as training set to predict the year 3.

| Mea<br>n | M1<br>($S+T+L$) | | M2<br>($S+T+g$) | | M3<br>($S+T+g+g\times S$) | | M4<br>($S+T+g+T\times S$) | | M5<br>($S+T+g+g\times T$) | | M6<br>($S+T+g+g\times S+T\times S+g\times T+g\times T\times S$) | |
| --- | --- | --- | --- | --- | --- | --- | --- | --- | --- | --- | --- | --- |
|  | PA | MS<br>E | PA | MS<br>E | PA | MS<br>E | PA | MSE | PA | MSE | PA | MSE |
| Y<sub>2</sub> | 0.41<br>0 | 2.34 | 0.41<br>4 | 2.28 | 0.65<br>9 | 1.90 | 0.41<br>3 | 2.28 | 0.39<br>3 | 2.37 | 0.669 | 2.30 |
| Y<sub>3</sub> | 0.17<br>4 | 7.30 | 0.27<br>3 | 7.26 | 0.42<br>8 | 7.11 | 0.27<br>3 | 7.27 | 0.26<br>6 | 7.30 | 0.393 | 7.45 |
| TRN Y<sub>2</sub> - TST Y<sub>3</sub> |  |  |  |  |  |  |  |  |  |  |  |  |
| Y<sub>3</sub> | 0.46<br>2 | 0.72 | 0.46<br>0 | 0.76 | 0.64<br>7 | 0.93 | 0.46<br>0 | 1.70 | 0.46<br>5 | 0.76 | 0.626 | 1.57 |
| TRN Y<sub>1</sub> & Y<sub>2</sub> - TST Y<sub>3</sub> |  |  |  |  |  |  |  |  |  |  |  |  |
| Y<sub>3</sub> | 0.41<br>8 | 5.22 | 0.44<br>1 | 4.94 | 0.61<br>9 | 5.02 | 0.43<br>9 | 4.22 | 0.43<br>9 | 5.12 | 0.634 | 5.11 |

For predicting year 3, the correlation reduced while the MSE increased compared to predicting year 2. Here, model M3 returns the best results (0.428, and MSE=7.11). The predictive ability of the model is high when using data from year 1 as a training set which could be helpful for identifying superior genotypes while shortening the breeding cycle.

In forward prediction scenario 2, utilizing the data from the second year to predict the third year, including the interaction between the marker SNPs and site (M3), returned a high correlation (0.647) and low MSE (0.93). Whereas the baseline model M1 without any genomic information showed a low predictive ability (0.46) and also a reduced MSE value (0.72).

The prediction scenario 3 combines data from the first and second years to predict the performance of genotypes in year 3. Here, the baseline model M1 returned to a correlation of 0.41 and a high MSE (5.22). The highest correlation (0.634) was obtained with model M6 for an MSE value of 5.11, while the lowest MSE (4.22) was produced with M4 with a lower predictive ability of 0.43.

## 4 DISCUSSION

In this study genomic prediction models considering different manners to account for genotype-by-environment interaction were evaluated under novel cross-validation schemes introduced to study predictive ability in perennial crops. In both species, the inclusion of the different interaction terms significantly reduced the unexplained residual variance. Also, the inclusion of interaction between harvest time and site captures the largest amount of phenotypic variability highlighting the importance of distinguishing when the phenotypes were collected.

Regarding the predictive ability of the models, the interactions helped to improve the results under the CV1 and CV2 terms while for CV0 and CV00 the main effect models performed better. The obtained results provide a comprehensive overview of advantages and limitations of using elaborated prediction models accounting for the genotype-by-environment interaction in different manners. Harvest time resulted to be a significant element to consider in interaction with site or location. Also, the proposed cross-validation schemes can be applied to other perennial species.

The GS models that account for the interaction between genes and the environmental factors related to site ***S***, and harvest time ***T*** could assist breeders to identify genotypes with optimum overall performance across site-harvest time combinations and specific performance of genotypes in these combinations. For example, studying annual crops, Roorkiwal et al. (2018) and Crossa et al. (2022) showed that the inclusion of the genotype-by-environment interaction outperforms the main effects models. In addition, a study on barley by Gillberg et al. (2019) demonstrated that incorporating the G×E interaction in genomic prediction models substantially improves the prediction accuracy compared to the models that ignore this term. Similar types of approaches could help to identify the environmental covariates and range of values that could be beneficial to improve the *Miscanthus* yield production.

### 4.1 Miscanthus sacchariflorus

#### 4.1.1 Variance components

The variance components were computed to estimate the percentage of variability captured by each one of the model terms of these six linear predictors. The most comprehensive model (M6), which includes all the main effects and interaction terms, showed the lowest residual variance. The main effect of harvest time ***T*** and the interaction between harvest time and site ***T***×***S*** explained higher variability than the other model terms because there is heterogeneity of variances associated to this factor. The phenotypic variance at the early stages (years 1 and 2) is lower due to the establishment of the crop than at the third year. In this case, the phenotypic variability exhibited in years 1, 2 and 3 were 13.3, 14.5, and 29.35, respectively. Previous studies have not considered interaction effects based on harvest time and site (Widener et al., 2024) in prediction models or more elaborated cross-validation schemes other than the conventional CV00 designed for annual crops (Jarquin et al., 2017). These authors considered the harvest time and site as unrelated environments within the same site; in our case we maintained a dependence structure between these. Here, we also introduce the forward prediction scheme for perennials as well.

Under M1, the site term ***S*** captured approximately 16% of the phenotypic variance. The inclusion of the main effect of the marker SNPs (**g**) addressed nearly 8% of the variability with M2 model. The variance component resulting from the interaction between these two factors captured around 15% in model M3 while the residual variance was reduced by 24%.

Furthermore, the other models, including interactions, also showed reduced residual variances compared to the main effect model M2. This indicates that the interactions captured a significant amount of phenotypic variability helping to better dissect the factors driving trait performance and consequently leverage this information to improve predictive ability. In this study, we did not attempt to model heterogeneity of variances mainly because the selection of outperforming genotypes is based on rankings only. These seemed not to be negatively affected by ignoring this assumption.

#### 4.1.2 Predictive ability

As expected, the performance of the prediction models depended on the different cross-validation schemes (Jarquin et al., 2021). In general, the baseline model without the marker data showed the lowest prediction accuracy. However, in some cases, their values were high, especially in those where the target was to predict already observed genotypes (CV2, CV0, and forward prediction). In general, the inclusion of marker SNPs increased the prediction accuracy, especially in the more challenging scenarios when phenotypic information of the testing genotypes is absent (CV1 and CV00). The effect of incorporating different interactions on predictive ability varied according to the different cross-validation schemes. Interestingly, in some cases, reduced MSE was accomplished for the same levels of predictive ability (e.g., M3 and M6 for CV1; M5 and M6 for CV2; M3 and M4 for CV0; M4 and M5 for CV00). The four k-fold cross-validation schemes (CV2, CV1, CV0, and CV00) have been implemented in several annual crops, such as wheat (Jarquin et al., 2017) and soybean (Persa et al., 2021). To our knowledge, this is the first time these four schemes are considered on perennial crops since their implementation is not straight forward. Extra care should be taken to avoid contradictory or impractical situations in perennial crops. For example, in *Miscanthus* breeding, predicting the performance of genotypes in Y_1_ using their phenotypic information from Y_2_ and/or Y_3_. The forward prediction has been implemented in annual crops for predicting trait performance within the same growing season using repeated measures (Momen et al., 2019). In addition, this prediction scheme has also been implemented on perennial tree species but for selecting offspring to be used as parents in future crosses (Lenz et al., 2020). To our knowledge forward prediction has not been implemented in perennial species such as *Miscanthus* for predicting trait performance of genotypes using data observed at early stages. Here, it is crucial to consider the repeated measures framework to leverage data previously collected.

The CV2 scheme showed a high prediction accuracy for all models because observing a genotype at any site provides good information to predict its performance in other sites. The **g**×***S*** interactions in M3 returned a prediction accuracy of 0.67, which represents an increase of 10% compared to model M2 (0.61). On the other hand, M6 showed a prediction accuracy of 0.66 while its corresponding MSE value was of 0.16, which represents a reduction of 80% with respect to M3 and 98% with respect to M2.

The CV1 scheme represents the prediction of untested genotypes in the observed environments (i.e., newly developed genotypes). Interestingly, for most of the models using genomic data, under this more challenging cross-validation scenario a higher prediction accuracy was accomplished compared to the CV2 scheme. This could be attributed to the availability of correlated site information. Considering the **g**×***S*** interaction in model M3 a slightly higher prediction accuracy (0.67) was obtained representing an increase of 3% compared to model M2 (0.65) and around 1% compared to model M6 (0.66). However, model M6 showed a significantly lower MSE value (0.02), which represents a reduction of 92% compared M3 (0.77). These results indicate that model M6 outperformed all the other considered models, returning a high accuracy and reduced MSE values. This model leverages the information captured by all the interaction terms, helping to borrow information among genotypes evaluated at different sites and harvest times.

Analyzing our results in CV2 and CV1, the results were benefited by the inclusion of different G×E interaction components such as the response of genotypes observed at different sites and times **g**×***S***. *Miscanthus* exhibited varied phenotypic variability across sites and at different harvest times. In the CV2 and CV1 schemes, models M3 and M6 outperformed other models in terms of prediction accuracy and MSE values. Thus, breeders should consider the mentioned interactions, especially the **g**×***S*** interaction to make informed decisions.

In our case, the CV0 scheme mimics the problem of predicting observed genotypes in unobserved sites across years. Here, models M2 and M5 showed the highest prediction accuracy (0.56), which represents an increase of 6% in comparison to models M3 and M4, both having the same accuracy (0.53). On the contrary, model M6 showed the lowest prediction accuracy (0.36) while M1, which does not include genomic information but only relies on phenotypic data observed in other sites and years, returned a higher correlation (0.43). The MSE values expressed by M2 (18.83) and M5 (17.76), respectively, represent an increase of 12% and 6%, compared to model M3 (16.8). However, there are no significant improvements in predictive ability neither in the reduction of MSE considering models that include interactions. Here, most of the genotypes to predict in a novel site were already observed at other sites. Hence, the information of already observed genotypes provides enough signal to predict trait performance at unobserved sites with high accuracy.

The CV00 scheme mimics the scenario evaluating new cultivars at new sites. Here, model M2 outperformed other models, showing a prediction accuracy of 0.59, which represents an increase of 18%, 2%, and 75% when compared to the model M3 (0.50), M5 (0.58), and M6 (0.34), respectively. With respect to MSE, model M1 showed the lowest MSE value (11.25); however, it also showed a negative predictive ability. This model lacks information to link genotypes in training and testing sets impeding the prediction of untested genotypes. Thus, in this case, the low MSE shown by this model becomes irrelevant, and we consider the predictive ability as the main criterion for selecting the best models. These results differ from Widener et al. (2024), where these authors accomplished higher prediction values (0.33∼0.91) for the *Msa* population, probably due to a different definition of the environment (site-by-location combination) implementing models that did not account by the harvest time either.

Overall, for the CV0 and CV00 schemes, the main effect model outperformed the interaction models. Thus, the most parsimonious models are preferable in these cases. Models having main effects performed better, or at least at the same level as those that included interaction effects, indicating that the main effects are adequate to capture the phenotypic variation and to make accurate predictions in these complex prediction scenarios. In our case, we took extra care to avoid using phenotypic information from future years to predict previous years or contamination when predicting future years in the same site under these four cross-validation schemes.

Therefore, we can infer that the impacts of including different interaction terms in the prediction models depend on the specific cross-validation scenarios that mimic the real problems faced by the breeders. However, accounting for the interactions can be beneficial in most scenarios for capturing phenotypic variations accomplishing sometimes more accurate predictions and helping to reduce the MSE. The main effects alone can also provide good prediction accuracy in some scenarios, but these might not be sufficient for capturing the genotype effects across sites at different harvest times. The decision could be based on the problem faced by the breeders and then using the appropriate cross-validation scheme to identifying the most accurate model for the specific prediction scenario.

#### 4.1.3 Forward prediction

Regarding the forward prediction for *M. sacchariflorus*, the second and third years were predicted using the first year’s observation data as a training set for scenario 1. The inclusion of **g** ×***S*** interactions in M3 model helped to return the highest predictive ability values (0.66 for Y_2_, and 0.57 for Y_3_). In addition, also the lowest MSE values were obtained with this model for these years, 11.31 and 31.00, respectively. In the forward prediction scenario 2, predicting the third year using the second year’s data as a training set, model M3 showed again the highest prediction accuracy (0.65), with also the lowest MSE value (16.25). Similar results were obtained for the forward prediction scenario 3 combining data from the first and the second year to predict the third year. Model M6 showed the highest prediction accuracy (0.66) and a MSE value of 30.53. However, M3 showing a similar prediction accuracy of 0.63 returned the lowest MSE (23.15) which represents a reduction of 24% with respect to M6.

Overall, predicting Y_3_ the forward prediction scenario 1 showed high predictive ability values comparable to those from scenarios 2 and 3 that required information from the second harvest as well. Thus, the prediction scenario 1 is convenient for breeders because it enables the identification of superior cultivars at early stages, shortening the selection process by two years. Hence, significantly accelerating the breeding cycle of *M. sacchariflorus* for commercial production.

### 4.2 Miscanthus sinensis

For *M. sinensis*, 1,870 individuals of 280 unique clones were evaluated at four locations around the world for two to three years.

#### 4.2.1 Variance components

For *M. sinensis*, the site term (***S***) captured 17.2% of the phenotypic variance in model M1, while marker **g** data included in model M2 captured 8.1%. The inclusion of interaction between them in model M3 (**g** ×***S***) captured 13% and the residual variance was reduced by 30% with respect to M2. Furthermore, the other interaction models also showed a lower residual variance compared to the main effect models, indicating that the inclusion of interactions helps to address the unexplained variability. Similar to the previous population, the main effect ***T*** and the interaction between the harvest time and site ***T***×***S*** explained higher variability than the other model terms due to a clear heterogeneity of variances across harvests. The average phenotypic variance of phenotypes at the early stages might be lower due to damage that occurred during the first year of establishment. The average phenotypic variance across the sites per year showed the lowest values in Y_1_ (0.64), then Y_2_ (1.26), and finally Y_3_ showed the highest variance (2.86).

#### 4.2.2 Predictive ability

Under the CV2 scheme, M6 the most comprehensive model, showed the highest prediction accuracy (0.68) equivalent to an increase of 8% and 30% with respect to M3 (0.63) and M2 (0.52) models, respectively. Also, it showed a low MSE value (0.62) which was reduced by 36% and 41% compared to M3 (0.98) and M2 (1.06), respectively.

For CV1, model M6 showed a high prediction accuracy of 0.48, which represents an increase of 14% compared to model M3 (0.42) and a 17% increase compared to model M2 (0.41). Another advantage of the M6 model is that it showed the lowest MSE value (0.81), representing a relative reduction by 22% with respect to M3 (1.04) and 25% for M2 (1.08).

For these reasons, for CV2 and CV1 cross-validation schemes M6 is considered superior to the others. Their terms, borrow more efficiently the information among genotypes at different sites and harvest times.

Under CV0, M4 and M5 interaction models showed the same prediction accuracy (0.13) with M4 being preferable by presenting a reduced MSE value (2.62). Compared to M2, these models slightly increased the predictive ability by 8% and 44%, and 160% with respect to model M6 (0.05). In addition, M4 reduced the MSE by 8% with respect to model M2 (2.82) and 4% for M5 (2.86) while compared to M6 it slightly increased this value by 1%. However, as mentioned M6 predictive ability is very low (0.05) hence the reduction in MSE becomes irrelevant under this model.

In CV00 similar patterns were shown for the interaction models M4 and M5 to those from CV0 where the same prediction accuracy (0.07) was obtained. It represents an increase of 16% compared to M2 (0.06), and 133% compared to M3 and M6 (0.03). In addition, model M4 showed a slightly lower MSE (2.55) representing a reduction of 9% in comparison to model M2 and M5, both having an MSE of 2.81. Similarly, a reduction of 8% and 2% was observed in comparison to M3 (2.77) and M6 (2.59), respectively. These results differ from Widener et al. (2024), where they have a prediction value of (0.10∼0.46) for the *Msi* population. These differences in the levels of predictive ability are mostly due to the adopted definition of environments in both studies.

In *M. sinensis*, for all cross-validation scenarios, in general the interaction models outperformed the models without interaction as the interactions capture the important patterns or relationships of the data, which leads to significant improvements in the prediction accuracy.

#### 4.2.3 Forward prediction

For the *M. sinensis* species, using the first year’s data as the training set, models M6 (∼0.67) and M3 (∼0.43) showed the highest prediction accuracy for predicting Y_2_ and Y_3_, respectively. In addition, model M3 showed the lowest MSE value for both years and a comparable predictive ability to M6 (∼0.66). Hence M3 model is considered superior to the others when using data from Y_1_ for model calibration to predict future years. Similar results were obtained in the forward prediction scenarios 2 and 3 for predicting Y_3_ using Y_1_ and combined Y_1_ and Y_2_ for model calibration. Here, consistently across both years, M3 and M6 models outperformed the others, showing the highest prediction accuracies for these years ∼0.65 and ∼0.63, respectively.

Overall, dealing with forward prediction in *M. sinensis,* models M3 and M6 including the **g**×***S*** interaction term showed the highest prediction accuracy. This shed light about genotypic responses being heavily affected by the interaction between genotypes and locations. These results also suggest that the first year’s observation data can serve as an effective training set for making predictions of the subsequent years. This implies that having only information on the first-year observation dataset could be reliable for making predictions for subsequent years. This approach would significantly reduce phenotyping costs, field evaluation of multiple genotypes across the locations for years optimizing the resource allocation.

## 5 CONCLUSIONS

The integration of genomic selection in *Miscanthus* breeding programs could be beneficial to help breeders meet the demand for biomass yield for biofuel and bioproduct production. GS holds the potential to accelerate the breeding cycle of this perennial crop, which requires up to three years of field testing to deliver reliable phenotypic information to make informed decisions. For *M. sacchariflorus* and *M. sinensis*, the interaction models performed better than the main effect models in most of the cross-validation scenarios (k-fold based and forward prediction) highlighting the importance of considering the inclusion of different types of G×E interactions in prediction models.

In addition, predicting biomass yield for subsequent breeding years using historical data (forward prediction) returned a high accuracy for selecting superior cultivars. Indicating that an important portion of the genetic pulses of the genotypes measured at early stages remain in the next years of evaluation. This could be helpful for significantly shortening *Miscanthus* breeding cycle. These findings indicate that the proposed models, including different genotype-by-environment interactions, can be beneficial for *Miscanthus* breeding, utilizing the first-year data for conducting predictions for subsequent years, helping breeders to make informed decisions for selecting top candidates ahead of time.

## ACKNOWLEDGMENTS

This work was funded by the DOE Center for Advanced Bioenergy and Bioproducts Innovation (U.S. Department of Energy, Office of Science, Office of Biological and Environmental Research under Award Number DE-SC0018420). Any opinions, findings, and conclusions or recommendations expressed in this publication are those of the authors and do not necessarily reflect the views of the U.S. Department of Energy.

## DATA AVAILABILITY

The full prediction pipeline can be found in https://doi.org/10.6084/m9.figshare.31768948

## SUPPLEMENTARY DATA

**Table S1.** Withing location-by-harvest average predictive ability comparison of six genomic selection prediction models across four cross-validation scenarios-A (CV2), B (CV1), C (CV0), and D (CV00) for biomass yield of *Miscanthus sacchariflorus* (MSA). M1 (*S*+*T*+*L*) site, time, and line main effects; M2 (*S*+*T*+g) also incorporating genomic information; M3 (*S*+*T*+g+g×*S*), accounts for by the genotype-by-site interaction; M4 (*S*+*T*+g+*T*×*S*) includes the site-by-time interaction; M5 (*S*+*T*+g+g×*T*) modeling the genotype-by-time interaction; M6 (*S*+*T*+g+g×*S*+*T*×*S*+g×*T*+g×*T*×*S*) adding all interactions including genotype-by-time-by-site. Locations: Aarhus University (AU), Hokkaido University (HU), University of Illinois (UI), Kangwon National University (KNU), and Zhejiang University (ZJU).

| CV2 |  |  |  |  |  |  |  |
| --- | --- | --- | --- | --- | --- | --- | --- |
| | | M1:<br>$S+T+L$ | M2:<br>$S+T+g$ | M3:<br>$S+T+g+g \times S$ | M4:<br>$S+T+g+T \times S$ | M5:<br>$S+T+g+g \times T$ | M6:<br>$S+T+g+g \times S+T \times S+g \times T+g \times T \times S$ |
| Env | SS | Mean | Mean | Mean | Mean | Mean | Mean |
| AU.Y<sub>1</sub> | 44<br>6 | 0.315 | 0.332 | 0.265 | 0.329 | 0.330 | 0.164 |
| AU.Y<sub>2</sub> | 44<br>3 | 0.299 | 0.322 | 0.250 | 0.319 | 0.319 | 0.210 |
| AU.Y<sub>3</sub> | 44<br>6 | 0.373 | 0.362 | 0.355 | 0.362 | 0.366 | 0.412 |
| HU.Y<sub>1</sub> | 35<br>6 | 0.323 | 0.360 | 0.372 | 0.357 | 0.357 | 0.303 |
| HU.Y<sub>2</sub> | 33<br>8 | 0.500 | 0.579 | 0.593 | 0.574 | 0.578 | 0.578 |
| HU.Y<sub>3</sub> | 33<br>8 | 0.521 | 0.605 | 0.599 | 0.598 | 0.603 | 0.590 |
| UI.Y<sub>1</sub> | 18<br>9 | 0.663 | 0.885 | 0.918 | 0.874 | 0.881 | 0.918 |
| UI.Y<sub>2</sub> | 18<br>9 | 0.638 | 0.857 | 0.894 | 0.846 | 0.852 | 0.897 |
| UI.Y<sub>3</sub> | 18<br>9 | 0.662 | 0.895 | 0.932 | 0.880 | 0.900 | 0.932 |
| KNU.Y<sub>1</sub> | 39<br>6 | 0.547 | 0.620 | 0.616 | 0.614 | 0.617 | 0.604 |
| KNU.Y<sub>2</sub> | 39<br>5 | 0.345 | 0.413 | 0.424 | 0.411 | 0.415 | 0.418 |
| KNU.Y<sub>3</sub> | 39<br>6 | 0.05 | 0.089 | 0.100 | 0.085 | 0.089 | 0.044 |
| ZJU.Y<sub>1</sub> | 40<br>7 | 0.505 | 0.691 | 0.752 | 0.683 | 0.690 | 0.765 |
| ZJU.Y<sub>2</sub> | 41<br>1 | 0.423 | 0.606 | 0.712 | 0.600 | 0.609 | 0.750 |
| ZJU.Y<sub>3</sub> | 41<br>1 | 0.499 | 0.691 | 0.760 | 0.681 | 0.693 | 0.758 |

S1.B
| CV1 |  |  |  |  |  |  |  |
| --- | --- | --- | --- | --- | --- | --- | --- |
| | | **M1:**<br>$S+T+L$ | **M2:**<br>$S+T+g$ | **M3:**<br>$S+T+g+g \times S$ | **M4:**<br>$S+T+g+T \times S$ | **M5:**<br>$S+T+g+g \times T$ | **M6:**<br>$S+T+g+g \times S+T \times S+g \times T+g \times T \times S$ |
| Env | SS | Mean | Mean | Mean | Mean | Mean | Mean |
| AU.Y<sub>1</sub> | 44<br>6 | 0.010 | 0.332 | 0.345 | 0.331 | 0.331 | 0.226 |
| AU.Y<sub>2</sub> | 44<br>3 | 0.005 | 0.326 | 0.335 | 0.324 | 0.323 | 0.307 |
| AU.Y<sub>3</sub> | 44<br>6 | -0.006 | 0.329 | 0.366 | 0.327 | 0.334 | 0.430 |
| HU.Y<sub>1</sub> | 35<br>6 | -0.033 | 0.362 | 0.365 | 0.361 | 0.359 | 0.290 |
| HU.Y<sub>2</sub> | 33<br>8 | -0.066 | 0.584 | 0.586 | 0.582 | 0.585 | 0.575 |
| HU.Y<sub>3</sub> | 33<br>8 | -0.083 | 0.608 | 0.595 | 0.605 | 0.605 | 0.588 |
| UI.Y<sub>1</sub> | 18<br>9 | -0.117 | 0.912 | 0.915 | 0.912 | 0.912 | 0.916 |
| UI.Y<sub>2</sub> | 18<br>9 | -0.110 | 0.881 | 0.889 | 0.881 | 0.881 | 0.892 |
| UI.Y<sub>3</sub> | 18<br>9 | -0.112 | 0.924 | 0.929 | 0.921 | 0.923 | 0.928 |
| KNU.Y<sub>1</sub> | 39<br>6 | -0.078 | 0.616 | 0.608 | 0.613 | 0.614 | 0.598 |
| KNU.Y<sub>2</sub> | 39<br>5 | -0.068 | 0.406 | 0.414 | 0.406 | 0.409 | 0.407 |
| KNU.Y<sub>3</sub> | 39<br>6 | -0.065 | 0.102 | 0.106 | 0.100 | 0.100 | 0.047 |
| ZJU.Y<sub>1</sub> | 40<br>7 | -0.041 | 0.734 | 0.753 | 0.734 | 0.736 | 0.766 |
| ZJU.Y<sub>2</sub> | 41<br>1 | -0.044 | 0.658 | 0.717 | 0.660 | 0.666 | 0.753 |
| ZJU.Y<sub>3</sub> | 41<br>1 | -0.049 | 0.742 | 0.765 | 0.743 | 0.739 | 0.760 |

**S1.C**
| CV0 |  |  |  |  |  |  |  |
| --- | --- | --- | --- | --- | --- | --- | --- |
|  |  | **M1:**<br>*S+T+*<br>*L* | **M2:**<br>*S+T+*<br>*g* | **M3:**<br>*S+T+g+g×*<br>*S* | **M4:**<br>*S+T+g+T×*<br>*S* | **M5:**<br>*S+T+g+g×*<br>*T* | **M6:**<br>*S+T+g+g×S+T×S+g×T+g×T*<br>*×S* |
| Env | SS | Mean | Mean | Mean | Mean | Mean | Mean |
| AU.Y<sub>1</sub> | 44<br>6 | 0.313 | 0.327 | 0.333 | 0.323 | 0.327 | 0.308 |
| AU.Y<sub>2</sub> | 44<br>3 | 0.297 | 0.318 | 0.325 | 0.311 | 0.317 | 0.300 |
| AU.Y<sub>3</sub> | 44<br>6 | 0.371 | 0.361 | 0.370 | 0.358 | 0.364 | 0.357 |
| HU.Y<sub>1</sub> | 35<br>6 | 0.316 | 0.333 | 0.285 | 0.349 | 0.345 | 0.220 |
| HU.Y<sub>2</sub> | 33<br>8 | 0.485 | 0.522 | 0.437 | 0.549 | 0.529 | 0.312 |
| HU.Y<sub>3</sub> | 33<br>8 | 0.498 | 0.558 | 0.460 | 0.571 | 0.569 | 0.382 |
| UI.Y<sub>1</sub> | 18<br>9 | 0.658 | 0.855 | 0.827 | 0.818 | 0.849 | 0.596 |
| UI.Y<sub>2</sub> | 18<br>9 | 0.635 | 0.823 | 0.806 | 0.791 | 0.817 | 0.554 |
| UI.Y<sub>3</sub> | 18<br>9 | 0.656 | 0.860 | 0.835 | 0.840 | 0.868 | 0.616 |
| KNU.Y<sub>1</sub> | 39<br>6 | 0.531 | 0.591 | 0.520 | 0.566 | 0.545 | 0.160 |
| KNU.Y<sub>2</sub> | 39<br>5 | 0.339 | 0.389 | 0.342 | 0.378 | 0.381 | 0.211 |
| KNU.Y<sub>3</sub> | 39<br>6 | 0.052 | 0.083 | 0.081 | 0.078 | 0.096 | 0.079 |
| ZJU.Y<sub>1</sub> | 40<br>7 | 0.487 | 0.638 | 0.608 | 0.618 | 0.639 | 0.435 |
| ZJU.Y<sub>2</sub> | 41<br>1 | 0.403 | 0.539 | 0.516 | 0.532 | 0.538 | 0.416 |
| ZJU.Y<sub>3</sub> | 41<br>1 | 0.477 | 0.633 | 0.608 | 0.619 | 0.638 | 0.510 |

**S1.D**
| CV00 |  |  |  |  |  |  |  |
| --- | --- | --- | --- | --- | --- | --- | --- |
|  |  | **M1:**<br>*S+T+*<br>*L* | **M2:**<br>*S+T+*<br>*g* | **M3:**<br>*S+T+g+g×*<br>*S* | **M4:**<br>*S+T+g+T×*<br>*S* | **M5:**<br>*S+T+g+g×*<br>*T* | **M6:**<br>*S+T+g+g×S+T×S+g×T+g×T*<br>*×S* |
| Env | SS | Mean | Mean | Mean | Mean | Mean | Mean |
| AU.Y<sub>1</sub> | 44<br>6 | 0.004 | 0.329 | 0.328 | 0.322 | 0.328 | 0.272 |
| AU.Y<sub>2</sub> | 44<br>3 | 0.001 | 0.323 | 0.323 | 0.318 | 0.320 | 0.301 |
| AU.Y<sub>3</sub> | 44<br>6 | 0.005 | 0.323 | 0.330 | 0.321 | 0.328 | 0.308 |
| HU.Y<sub>1</sub> | 35<br>6 | -<br>0.007 | 0.334 | 0.267 | 0.341 | 0.349 | 0.234 |
| HU.Y<sub>2</sub> | 33<br>8 | -<br>0.008 | 0.543 | 0.427 | 0.566 | 0.549 | 0.337 |
| HU.Y<sub>3</sub> | 33<br>8 | -<br>0.015 | 0.567 | 0.452 | 0.596 | 0.578 | 0.418 |
| UI.Y<sub>1</sub> | 18<br>9 | 0.045 | 0.887 | 0.831 | 0.878 | 0.875 | 0.625 |
| UI.Y<sub>2</sub> | 18<br>9 | 0.037 | 0.857 | 0.807 | 0.833 | 0.846 | 0.521 |
| UI.Y<sub>3</sub> | 18<br>9 | 0.031 | 0.898 | 0.841 | 0.881 | 0.897 | 0.608 |
| KNU.Y<sub>1</sub> | 39<br>6 | -<br>0.029 | 0.581 | 0.468 | 0.572 | 0.557 | 0.162 |
| KNU.Y<sub>2</sub> | 39<br>5 | -<br>0.030 | 0.378 | 0.302 | 0.381 | 0.382 | 0.173 |
| KNU.Y<sub>3</sub> | 39<br>6 | -<br>0.007 | 0.102 | 0.079 | 0.094 | 0.102 | 0.075 |
| ZJU.Y<sub>1</sub> | 40<br>7 | -<br>0.004 | 0.652 | 0.586 | 0.649 | 0.659 | 0.410 |
| ZJU.Y<sub>2</sub> | 41<br>1 | 0.010 | 0.555 | 0.498 | 0.561 | 0.564 | 0.370 |
| ZJU.Y<sub>3</sub> | 41<br>1 | 0.001 | 0.662 | 0.590 | 0.651 | 0.668 | 0.484 |

**Table S2.**
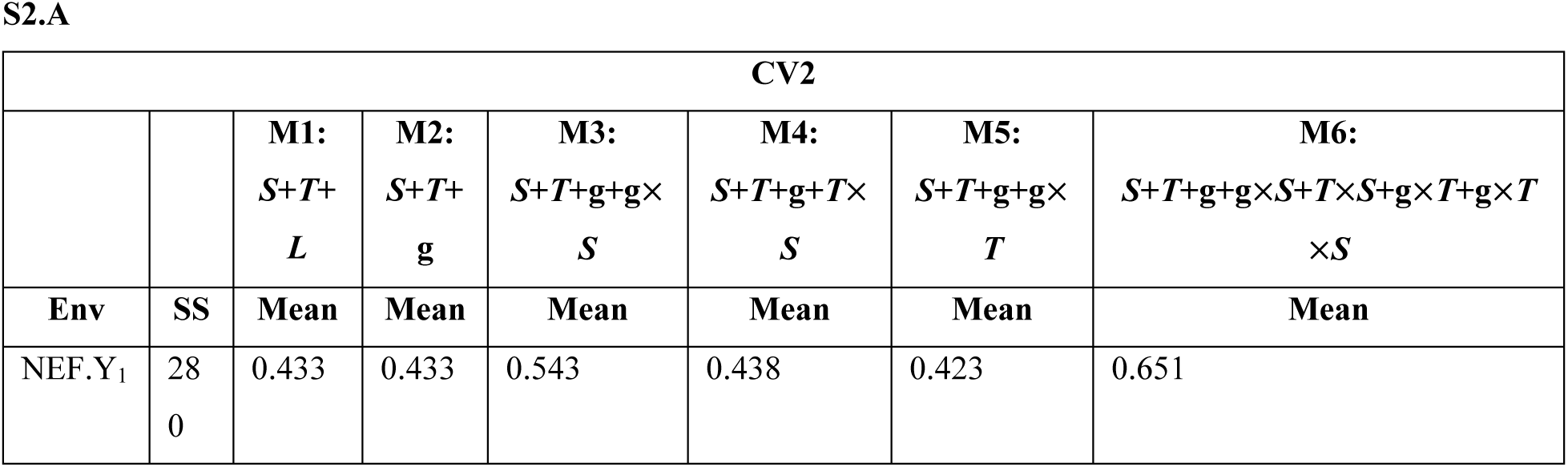
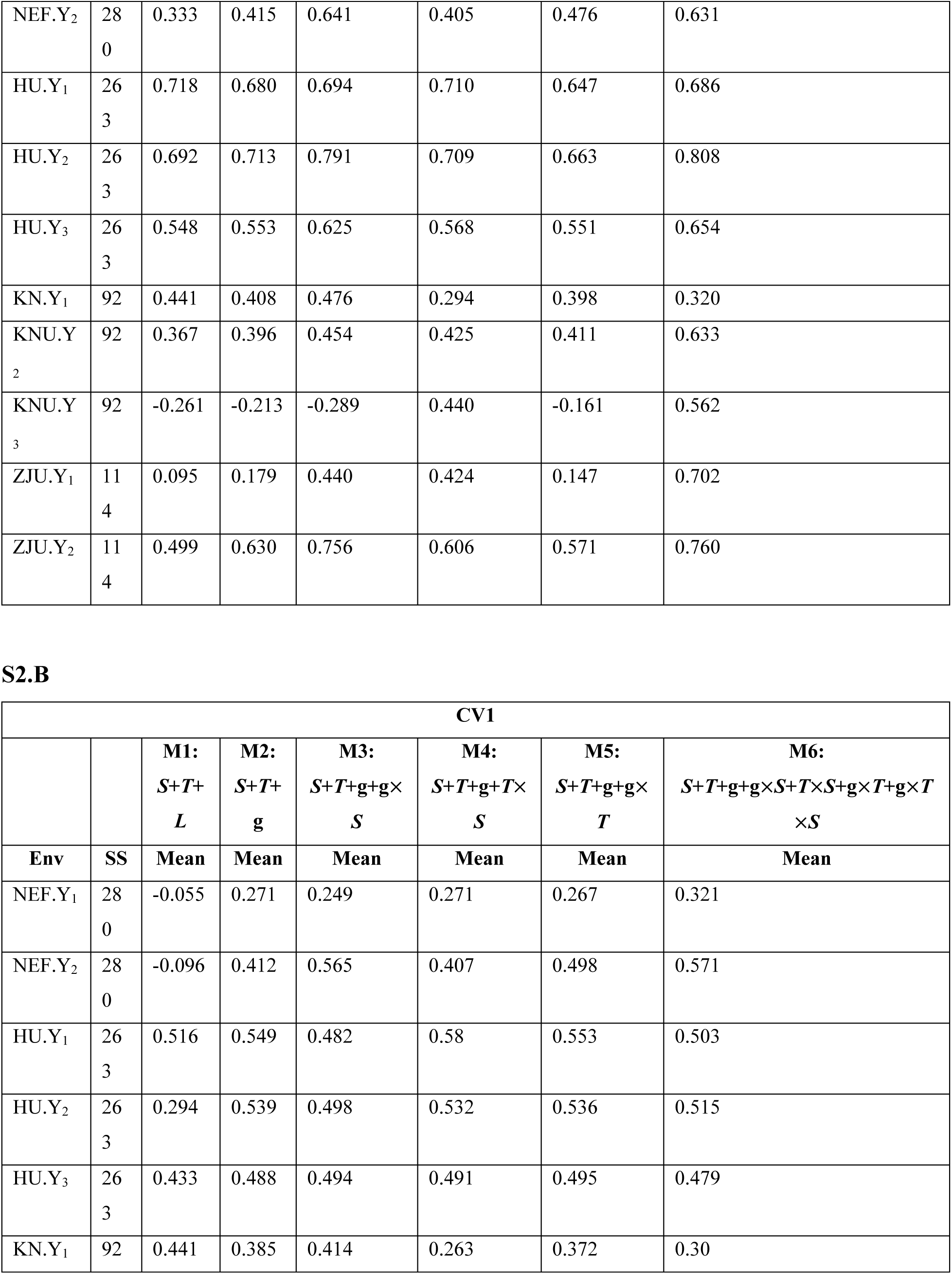
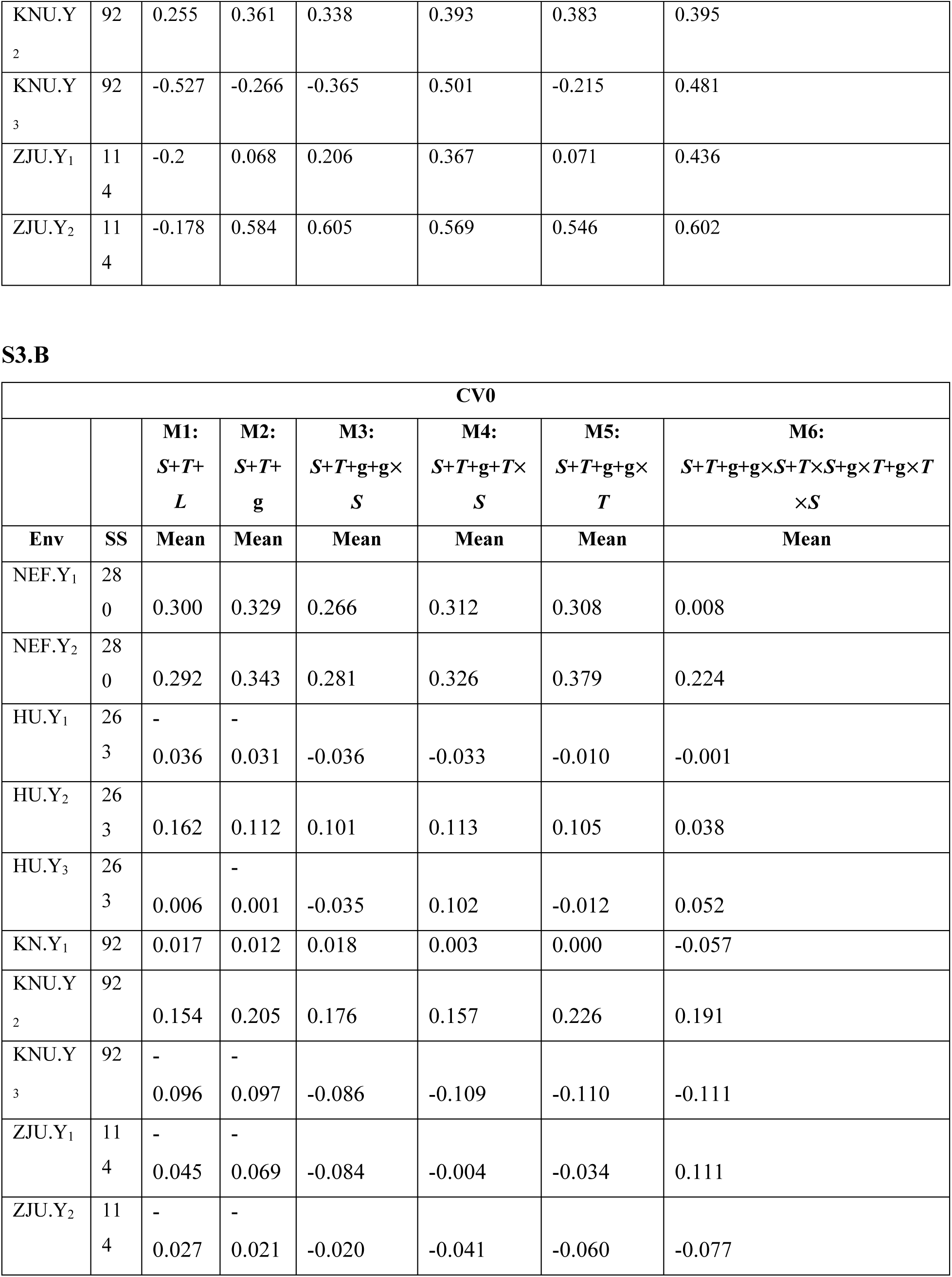
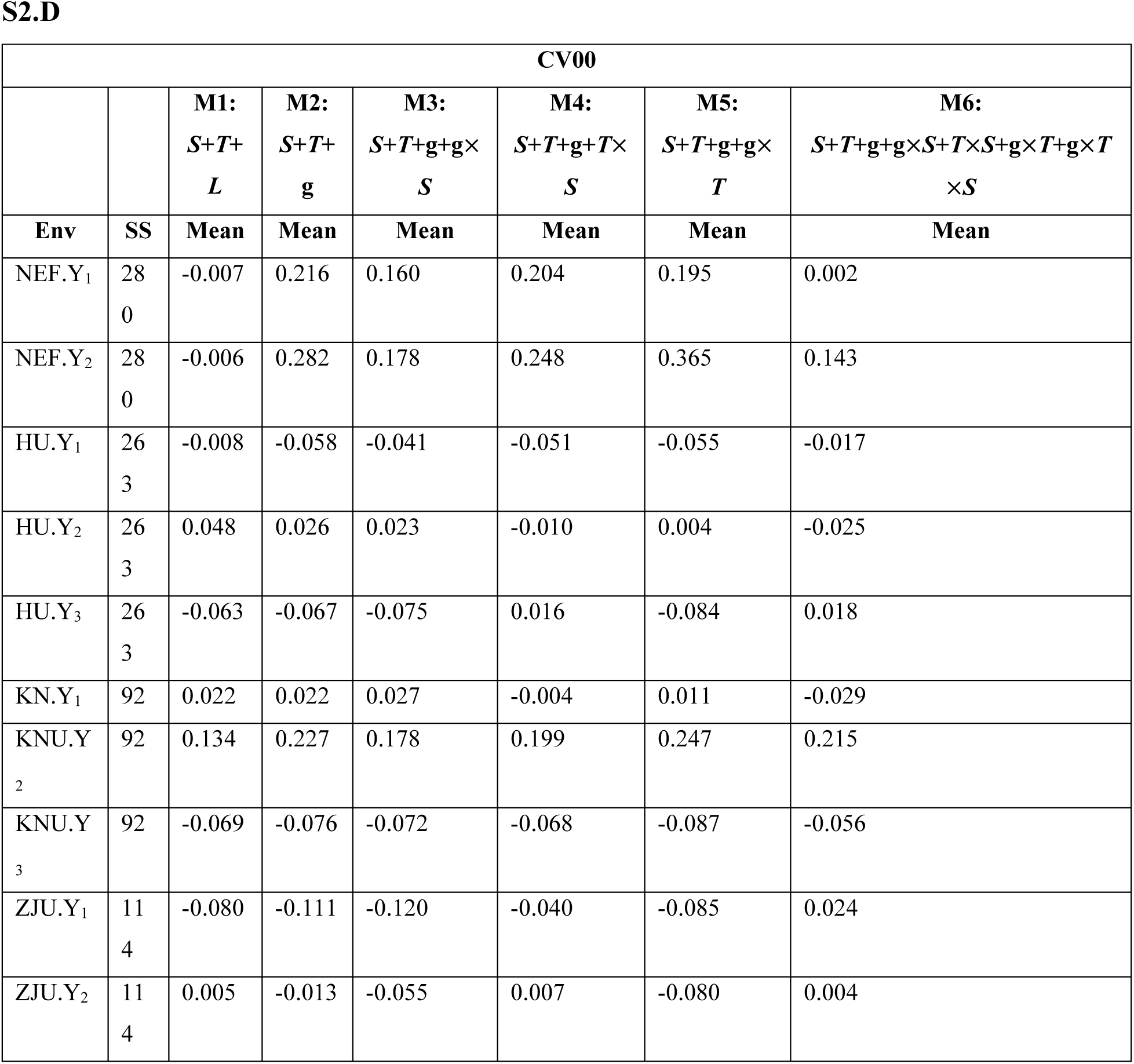
Withing location-by-harvest average predictive ability comparison of six genomic selection prediction models across four cross-validation scenarios-A (CV2), B (CV1), C (CV0), and D (CV00) for biomass yield of *Miscanthus sacchariflorus* (MSA). M1 (*S*+*T*+*L*) site, time, and line main effects; M2 (*S*+*T*+g) also incorporating genomic information; M3 (*S*+*T*+g+g×*S*), accounts for by the genotype-by-site interaction; M4 (*S*+*T*+g+*T*×*S*) includes the site-by-time interaction; M5 (*S*+*T*+g+g×*T*) modeling the genotype-by-time interaction; M6 (*S*+*T*+g+g×*S*+*T*×*S*+g×*T*+g×*T*×*S*) adding all interactions including genotype-by-time-by-site. Locations: New Energy Farms (NEF), Hokkaido University (HU), Kangwon National University (KNU), and Zhejiang University (ZJU).

## Abbreviations

CV: Cross-validation
G×E: Genotype-by-environment
GS: Genomic selection
*Msa*: *Miscanthus sacchariflorus*
*Msi*: *Miscanthus sinensis*
*Mxg*: *Miscanthus giganteus*
PA: Predictive Ability
MSE: Mean Square Error

